# CLPTM1L is a lipid scramblase involved in glycosylphosphatidylinositol biosynthesis

**DOI:** 10.1101/2021.07.12.451801

**Authors:** Yicheng Wang, Anant K. Menon, Yuta Maki, Yi-Shi Liu, Yugo Iwasaki, Morihisa Fujita, Paula A. Guerrero, Daniel Varón Silva, Peter H. Seeberger, Yoshiko Murakami, Taroh Kinoshita

## Abstract

Glycosylphosphatidylinositols (GPIs) are membrane anchors of many eukaryotic cell surface proteins. Biosynthesis of GPIs is initiated at the cytosolic face of the endoplasmic reticulum (ER) and the second intermediate, glucosaminyl-phosphatidylinositol (GlcN-PI), is translocated across the membrane to the lumenal face for later biosynthetic steps and attachment to proteins. The mechanism of the lumenal translocation of GlcN-PI is unclear. We report that Cleft lip and palate transmembrane protein 1-like protein (CLPTM1L), an ER membrane protein of unknown function, is a lipid scramblase involved in GPI biosynthesis. Purified CLPTM1L scrambles GlcN-PI, PI, and several other phospholipids in vitro. Knockout of *CLPTM1L* gene in mammalian cultured cells partially decreased GPI-anchored proteins due to impaired usage of GlcN-PI, suggesting a major role of CLPTM1L in lumenal translocation of GlcN-PI.

**One-Sentence Summary:** CLPTM1L translocates glucosaminyl-phosphatidylinositol across the membrane during glycosylphosphatidylinositol biosynthesis.

## Main Text

Glycosylphosphatidylinositol (GPI) anchoring is a post translational modification of many eukaryotic cell surface proteins, including more than 150 human proteins (*1*). GPI is a highly conserved complex glycolipid in eukaryotic organisms, with a core structure comprising EtNP-6Manα–2Manα–6Manα–4GlcNα–6Inositol-PL (where EtNP, Man, GlcN, and PL are ethanolamine phosphate, mannose, glucosamine, and phospholipid, respectively). Biosynthesis of GPI starting from phosphatidylinositol (PI) is a stepwise sequence of 11 reactions occurring on the endoplasmic reticulum (ER) membrane (Fig. 1A). The first biosynthetic step is transfer of *N*-acetylglucosamine (GlcNAc) from uridine-diphosphate (UDP)-GlcNAc to PI to generate GlcNAc-PI, which is then de-*N*-acetylated to generate GlcN-PI. These two enzymatic reactions occur on the cytosolic face (*2, 3*) and the resulting GlcN-PI is translocated across the ER membrane into the lumenal face, where later steps and attachment to proteins occur (Fig. 1A). Whereas enzymes that catalyze transfers of monosaccharides and other components to growing GPI have been molecularly cloned and characterized, the molecular basis of cytosol-to-lumen transmembrane translocation of GlcN-PI is unknown. It was suggested that GlcN-PI translocation is mediated by a scramblase without consumption of ATP (*4*), however its molecular identity is yet to be determined. Cytosol-to-lumen translocation across the ER membrane is a common process among lipid glycoconjugates involved in protein modifications, i.e., dolichol-pyrophosphate-heptasaccharide, dolichol-phosphate-mannose (DPM), and dolichol-phosphate-glucose are generated on the cytoplasmic face of the ER and then are translocated to the lumenal side. Their transmembrane translocations are also thought to be mediated by ER-resident scramblases, however, the underlying molecular mechanisms have not been clarified either (*5*–*8*). Many genes required for GPI biosynthesis have been identified by establishing GPI-deficient mutants from various cultured cell lines. Given that the entry of GlcN-PI into the ER lumen is an essential step for GPI assembly, it is surprising that previous intensive forward genetic screens all failed to identify GlcN-PI scramblase. It seems possible that more than one protein works redundantly to scramble GlcN-PI or that the scramblase is essential for cell survival for its other role.

**Fig. 1.**
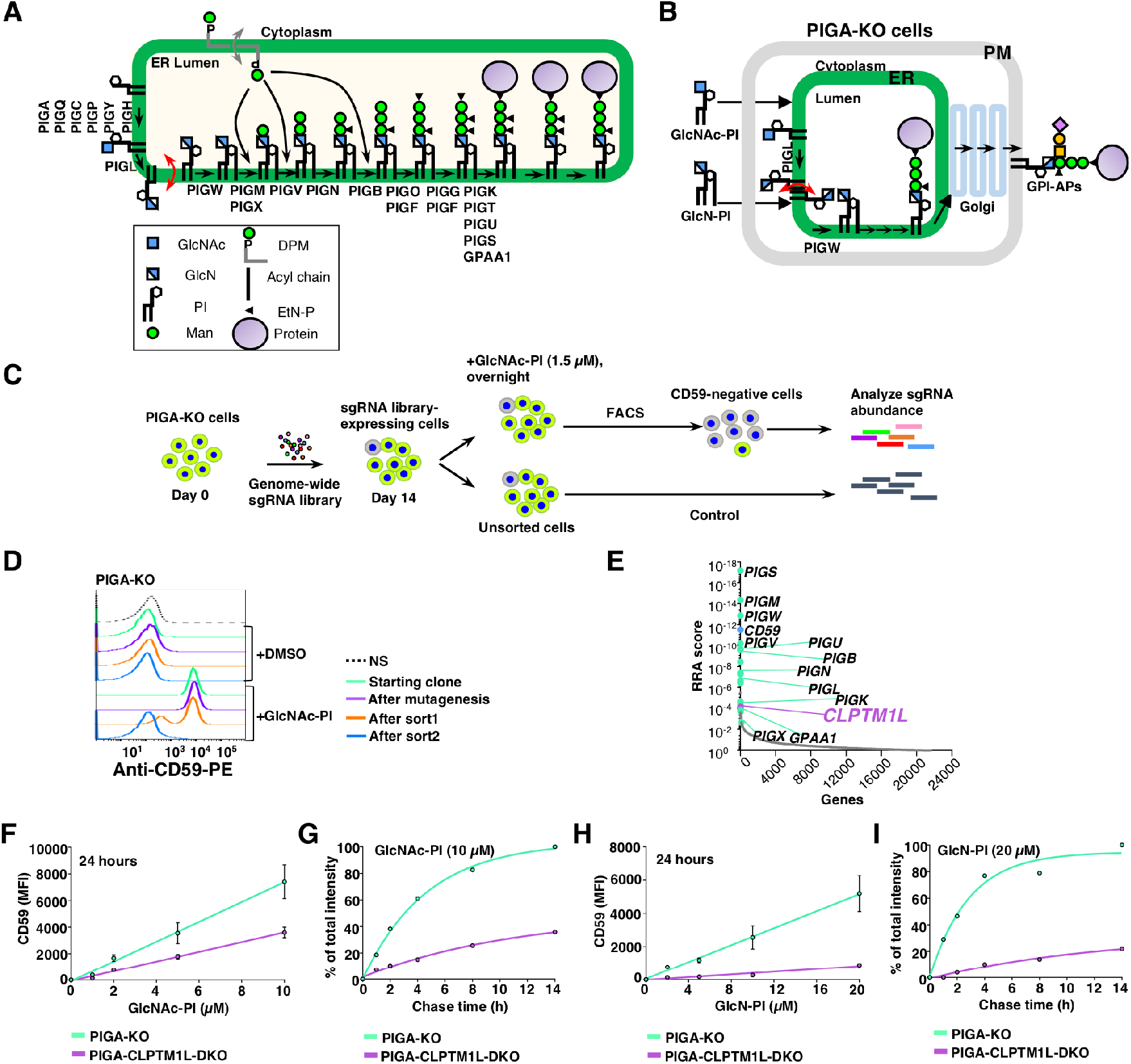
A genetic screen for components of the GPI biosynthetic pathway identified *CLPTM1L*. (**A**) Schematic of the GPI biosynthetic pathway in the ER. The GlcN-PI scrambling step is indicated in red, and DPM scrambling is indicated in gray. Genes essential for GPI biosynthesis and GPI anchoring to protein are listed. (**B**) Schematic of the restoration of GPI biosynthesis in PIGA-KO cells in the presence of chemically synthesized GlcNAc-PI or GlcN-PI. (**C**) Scheme depicting a FACS-based genome-wide CRISPR screen for genes involved in efficient GlcNAc-PI utilization for GPI biosynthesis using PIGA-KO HEK293 cells. (**D**) Loss of an ability to restore GPI-AP using exogenous GlcNAc-PI by PIGA-KO HEK293 cells after the FACS-based genome-wide CRISPR screen. Starting clone, cells after mutagenesis, sort1 cells, and sort2 cells incubated with DMSO or 2 µM GlcNAc-PI overnight were stained with anti-CD59 mAb and analyzed by flow cytometry. NS, cells not stained with 1^st^ antibody. (**E**) Gene scores in unsorted versus sorted PIGA-KO cells. Known GPI biosynthetic pathway genes are shown in green with some of their names, *CD59* is shown in blue, and *CLPTM1L* is shown in purple. (**F** and **H**) Restoration of CD59 expression in PIGA-KO and CLPTM1L-PIGA-DKO cells in the presence of various concentrations of GlcNAc-PI (F) or GlcN-PI (H) for 24 hours. Results are from two independent experiments. (**G** and **I**) Time course of CD59 expression in PIGA-KO and CLPTM1L-PIGA-DKO cells pre-incubated with GlcNAc-PI (G) or GlcN-PI (I) for 2 hours. Data points represent the average values of two independent experiments.

Aiming to identify more genes, including the GlcN-PI scramblase, that are involved in biosynthesis and trafficking of GPI-anchored proteins (GPI-APs), we took advantage of our recent finding that chemically synthesized GlcNAc-PI and GlcN-PI are able to rescue GPI-AP generation in PIGA-knockout (KO) cells that do not make GlcNAc-PI due to a defect in the first step in GPI biosynthesis (Fig. 1B) (Paula A. Guerrero et al., manuscript under review). We established a genome-wide CRISPR-Cas9 KO screen using PIGA-KO HEK293 cells as the parent cells. PIGA-KO cells were transduced with a genome-wide single guide RNA (sgRNA) library and maintained for two weeks, followed by culturing with GlcNAc-PI at a low concentration overnight (*9*). Mutant cells defective in restoring CD59, a GPI-AP marker, were collected by cell sorting and enriched sgRNA sequences determined (Fig. 1, C and D). As expected, most of previously known genes involved in GPI biosynthetic pathway were highly enriched in the sorted CD59-deficient cells. Notably, Cleft lip and palate transmembrane protein 1-like (*CLPTM1L*) of unknown molecular function was one of the top-ranking genes from this screen (Fig. 1E). To confirm the screening result, CLPTM1L-PIGA-double KO (DKO) HEK293 cells were generated (fig. S1, A and B) and their abilities to restore GPI-APs after incubation with GlcNAc-PI and GlcN-PI were assessed. GlcNAc-PI was 2-fold less effective in promoting CD59 synthesis in DKO vs PIGA-KO cells when measured at a single time point (Fig. 1F) and 3.3-fold less effective when assessed in a pulse-chase experiment (Fig. 1G). Restoration efficiency with GlcN-PI was even more strongly affected by CLPTM1L KO, being 5.1-fold and 10.6-fold less efficient in the two types of assays used (Fig. 1, H and I). Therefore, CLPTM1L is required for efficient generation of GPI-APs from GlcN-PI.

CLPTM1L, also known as cisplatin resistance-related protein 9, was originally identified as a gene associated with cisplatin-induced apoptosis of carcinoma cells and is a member of the CLPTM1 family (*10*). Consistent with involvement in GPI biosynthesis, CLPTM1L is ubiquitously expressed in human tissues and organs similar to PIGA (fig. S2, A and B). Both plasma membrane and intracellular localizations of CLPTM1L were observed in pancreatic adenocarcinoma cells and antibody-producing plasma cells, which express high levels of this protein (*11*–*13*). In human HEK293 cells, a non-carcinoma cell line, endogenous CLPTM1L protein detected by antibody, whose specificity was verified by using CLPTM1L-KO HEK293 cells (fig. S1A), colocalized with an ER membrane protein UBE2J1 (Fig. 2A). The *N*-glycans of CLPTM1L were sensitive to Endoglycosidase H treatment, consistent with ER localization of the protein (fig. S2C). Overexpressed monomeric enhanced green fluorescent protein (mEGFP) tagged CLPTM1L (CLPTM1L-mEGFP) failed to target to the cell surface of HEK293 cells (fig. S2D), further supporting the ER localization of CLPTM1L in these cells. Whereas in carcinoma cells, a fraction of CLPTM1L might exit the ER when it is expressed at high levels, our results indicate that CLPTM1L acts at the level of the ER in regulating GlcN(Ac)-PI-mediated restoration of GPI biosynthesis in PIGA-KO HEK293 cells.

**Fig. 2.**
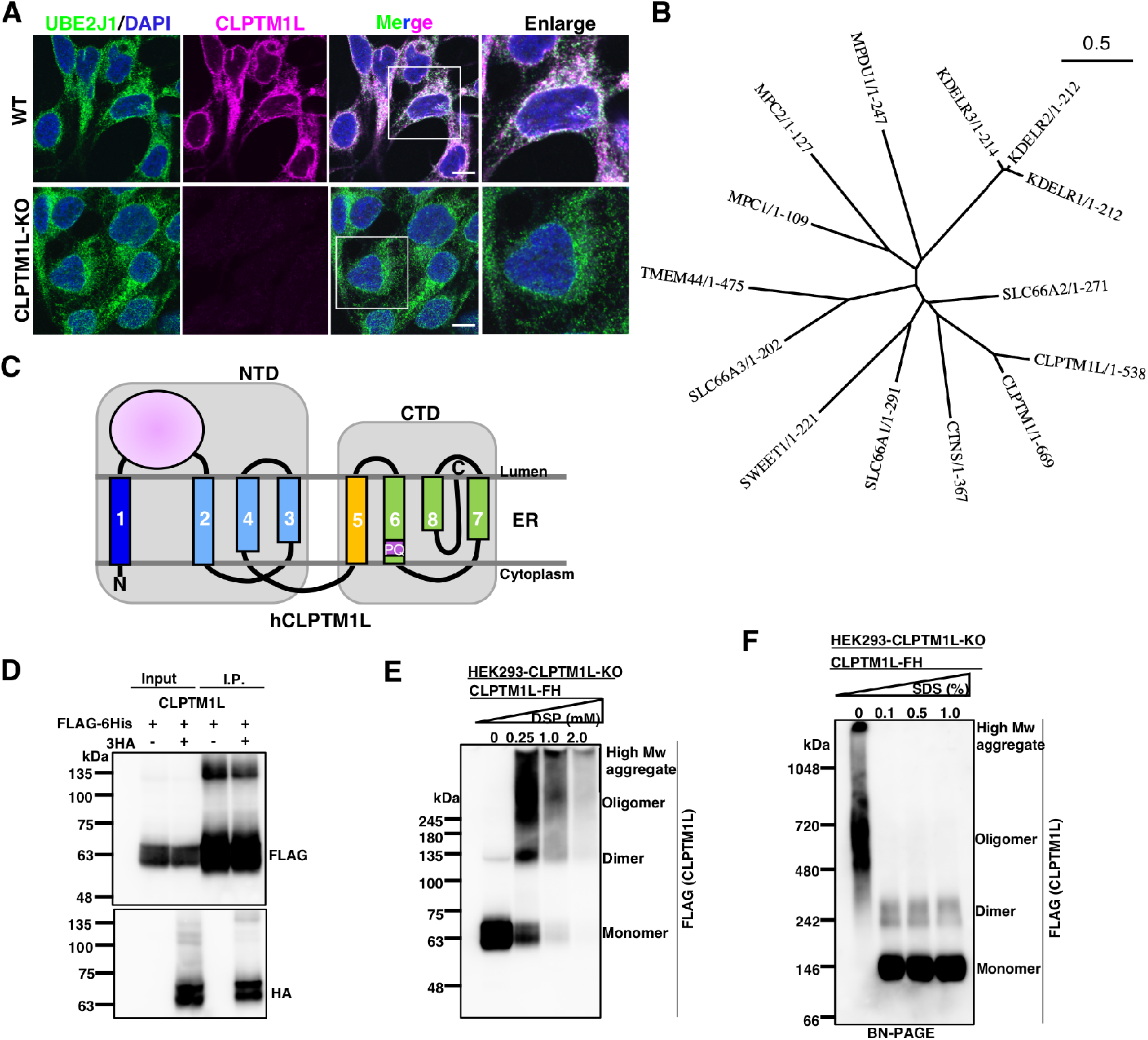
CLPTM1L is an ER scramblase candidate for GPI biosynthetic pathway. (**A**) Immunofluorescence detection of endogenous CLPTM1L (magenta) and the ER membrane protein marker UBE2J1 (green) in wild-type (WT) and CLPTM1L-KO HEK293 cells. Scale bar, 10 μm. (**B**) The human PQ-loop family tree. (**C**) Topology model of human CLPTM1L in the ER membrane. NTD, N-terminal domain; CTD, C-terminal domain. The TM domains are indicated by numbers. The PQ motif within TM6 is indicated. (**D**) Empty vector or CLPTM1L-3HA was co-expressed with CLPTM1L-FLAG-6His (CLPTM1L-FH) in CLPTM1L-KO HEK293 cells. Immunoprecipitates with anti-FLAG antibody resins were separated by SDS/PAGE and analyzed by western blotting with anti-FLAG and anti-HA antibodies. (**E**) CLPTM1L-KO HEK293 cells expressing CLPTM1L-FH were incubated with various concentrations of a chemical cross-linker DSP. Cell lysates were analyzed by western blotting. (**F**) Lysates from CLPTM1L-KO HEK293 cells expressing CLPTM1L-FH were treated with various concentrations of SDS. Proteins were separated by blue-native gel electrophoresis and were analyzed by western blotting.

Various algorithms predict 6-8 transmembrane (TM) domains in human CLPTM1L (fig. S3A). To verify the membrane topology of CLPTM1L, we determined the orientation of the hydrophilic region between TM1 and TM2, and the C-terminus in HEK293 cells. The region between TM1 and TM2 was detected by a region-specific antibody when both plasma membrane and the ER membrane were permeabilized by Triton X-100 but not when only plasma membrane was permeabilized by digitonin (The latter treatment allowed detection of a cytosolic protein, UBE2G2.) (fig. S3B). Likewise, mEGFP fused to the C-terminus of CLPTM1L and the ER lumenal protein PDIA4 were stained with monoclonal antibodies (mAbs) only after permeabilization by Triton X-100 (fig. S3C). Based on these results, we propose that CLPTM1L contains 7 TM domains, or 8 TM domains and an additional half-TM domain and that the C-terminus and the region between TM1 and TM2 are lumenally oriented (fig. S3D). We then generated a structure model for human CLPTM1L by trRosetta (*14*), which used two SWEET/ PQ-loop (PF04193) family members from *Arabidopsis thaliana*, atSWEET13 and atSWEET2, as the template proteins (fig. S4, A and B). SWEETs are bidirectional sugar transporters (*15*), which might also transport other metabolites, like gibberellins (*16*). The PQ-loop family is a functionally diverse family of membrane proteins including CLPTM1L (Fig. 2B) (*17*). The PQ-loop family proteins usually have seven TM domains in a 1–3–2–4–5–7–6 helix arrangement (*18*). Despite lack of significant sequence similarity between CLPTM1L and atSWEET13 (fig. S4A), the transmembrane arrangement of TM2 to TM8 from the predicted CLPTM1L structure has similarity to atSWEET13 (Fig. 2C, and fig. S4, C and D). Although no PQ-loop protein has been demonstrated to be a lipid scramblase, ANY1, a PQ-loop protein antagonizing yeast phospholipid flippase, was predicted as a lipid scramblase candidate (*19*). The PQ-loop family belongs to the transporter-opsin-G protein-coupled receptor (TOP) superfamily. Some TOP family proteins such as the rhodopsin-like class-A G protein-coupled receptors have lipid scramblase activity (*20, 21*). Furthermore, CLPTM1L, similar to many scramblases, appeared to make homodimers or bigger oligomers on analysis by co-immunoprecipitation, chemical cross-linking and blue-native PAGE (Fig. 2, D to F), supporting the idea that CLPTM1L may have a lipid scramblase activity.

To test whether CLPTM1L can scramble GlcN-PI across the membrane (Fig. 3A), we measured the scramblase activity of CLPTM1L in vitro. Bacterial PI-specific phospholipase C (PI-PLC) has been previously used as a membrane-impermeable topology probe for PI and GlcN-PI (*2, 22, 23*), and a scramblase assay using PI-PLC has been described to test activities of other scramblases, including opsin and nhTMEM16, towards [^3^H]PI in reconstituted vesicles (*24*). In this assay system, since PI-PLC only accesses the outer leaflet of liposomes and because spontaneous translocation of [^3^H]GlcN-PI across the membrane bilayer is expected to be negligible in a short time, about 50% of [^3^H]GlcN-PI will be cleaved on adding PI-PLC. If the vesicles possess a GlcN-PI scramblase, then greater than 50% of [^3^H]GlcN-PI will be hydrolyzed, because molecules from the inner leaflet of the vesicles will be translocated across the bilayer in a short period of time (Fig. 3B). To test whether CLPTM1L can translocate GlcN-PI, we purified CLPTM1L-FLAG-6His protein (Fig. 3C) and [^3^H]GlcN-PI (Fig. 3D), and reconstituted them into liposomes consisting of nine parts of egg phosphatidylcholine (PC) and one part of bovine brain phosphatidylserine (*21, 25*). In some experiments, [^3^H]PI was included instead of [^3^H]GlcN-PI. In an initial time course study, we observed approximately 80 % hydrolysis of GlcN-PI in CLPTM1L-proteioliposomes and approximately 50% hydrolysis of [^3^H]GlcN-PI in protein-free liposomes within 10 minutes (Fig. 3E). Similar results were obtained using [^3^H]PI (Fig. 3F), indicating that CLPTM1L facilitates transbilayer movement of both lipids. Consistent with a previous study using nhTMEM16 and opsin (*24*), hydrolysis of GlcN-PI or PI by PI-PLC in CLPTM1L-containing proteoliposomes was a biphasic process (Fig. 3, E and F), with an initial rapid burst corresponding to hydrolysis of substrate in the outer leaflet followed by a slower phase in which molecules from the inner leaflet cross the membrane to the outer leaflet. To test whether scrambling of GlcN-PI and PI is CLPTM1L-dose dependent, we measured the scramblase activity by incubating CLPTM1L-proteoliposomes with different protein to phospholipid ratios (PPRs) for 10 minutes. Hydrolysis of GlcN-PI and PI increased from ∼50% to >80% with increased PPRs from 0-7 mg/mmol (Fig. 3G), similar to a previous study using nhTMEM16 and opsin (*24*). To address whether CLPTM1L can translocate other kinds of PLs, we tested nitrobenzoxadiazole (NBD)-labeled PLs. The membrane-impermeant reductant dithionite bleaches the fluorescence of NBD-PLs in the outer leaflet of the liposomes (*20, 21*), thus, approximately 50% bleaching is expected on adding dithionite to protein-free liposomes and nearly complete reduction of fluorescence is expected if all the vesicles contain a scramblase (Fig. 3H). We prepared NBD acyl-labeled-PI (NBD-PI) as a control lipid (fig. S5, A to D). As seen by the greater loss of fluorescence in dithionite-treated CLPTM1L-proteoliposomes versus protein-free liposomes, CLPTM1L scrambles NBD-PI, as well as NBD acyl-labeled PC (NBD-PC), and NBD head group-labeled phosphatidylethanolamine (NBD-PE) (Fig. 3I). Therefore, CLPTM1L exhibits scramblase activity towards GlcN-PI, PI, PC, and PE in vitro.

**Fig. 3.**
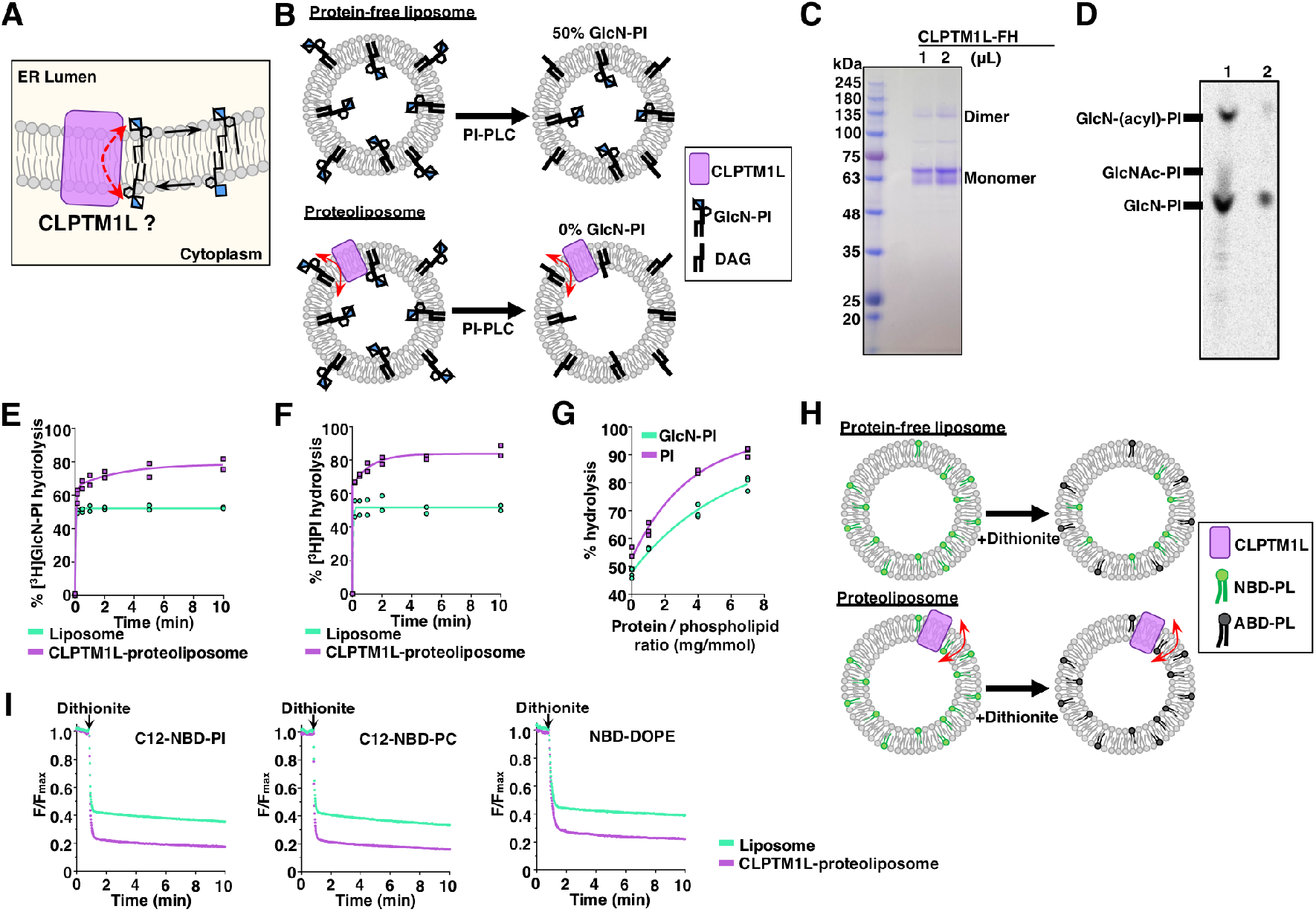
CLPTM1L scrambles GlcN-PI and various phospholipids in vitro. (**A**) Schematic of CLPTM1L-mediated ER lumenal translocation of GlcN-PI. (**B**) Schematic of PI-PLC based scramblase assay. PI-PLC hydrolyzes [^3^H]GlcN-PI in the outer leaflet of liposomes. (**C**) SDS-PAGE analysis of purified human CLPTM1L-FLAG-6His. (**D**) HPTLC analysis of early GPI precursors (lane 1) and purified [^3^H]GlcN-PI (lane 2). GPI precursors are labeled by UDP-[^3^H]GlcNAc in vitro using microsomes prepared from HEK293 cells (lane 1) and PIGU-defective CHO cells (lane 2). (**E** and **F**) Time course of [^3^H]GlcN-PI (E) or [^3^H]PI (F) hydrolysis by PI-PLC in CLPTM1L-proteoliposomes using protein to phospholipid ratio (PPR) of ∼7 mg/mmol. Protein-free liposomes were used as controls. Results are from two independent experiments. (**G**) PI-PLC mediated hydrolysis of [^3^H]GlcN-PI or [^3^H]PI in CLPTM1L-proteoliposomes for 10 min at different PPRs. Results are from three independent measurements using the same batch of [^3^H]GlcN-PI or [^3^H]PI containing liposomes. (**H**) Schematic of dithionite-based scramblase assay. NBD-PLs in the outer leaflet of liposomes are bleached to non-fluorescent 7-amino-2,1,3-benzoxadiazol-4-yl (ABD)-PLs by dithionite. (**I**) Substrate specificity of CLPTM1L towards acyl-NBD-PI (left), acyl-NBD-PC (middle), and N-NBD-DOPE (right). Scrambling of NBD-PLs in CLPTM1L-proteoliposomes using PPR of ∼1 mg/mmol. Similar results were observed from two independent experiments.

Results with CLPTM1L-PIGA-DKO HEK293 cells indicated that cytosol-to-lumen translocation of GlcN-PI is mediated redundantly by more than one scramblase and that CLPTM1L accounts for the major fraction of the translocation activity (Fig. 1, F-I). Next, we determined whether CLPTM1L is critical for steady state levels of GPI-APs by generating CLPTM1L-KO HEK293 cells. CLPTM1L-KO caused approximately a 30% reduction in cell surface CD59, which was rescued by expression of CLPTM1L-mEGFP (Fig. 4A, and fig. S6A). The partial deficiency (10% to 30% reduction) of CD59 was also observed in some but not all CLPTM1L-KO human cells derived from various cell lines (fig. S6, B and C). Therefore, it is likely that GlcN-PI scrambling is mediated by redundantly acting scramblases in various cell types. Overexpression of TMEM16K, a Ca^2+^ activated ER scramblase (*26*), failed to rescue GPI deficiency in CLPTM1L-KO cells (fig. S6, D and E). Although the other CLPTM1L homologous protein, CLPTM1, is likely a lipid scramblase, deficiency of GPI biosynthesis in CLPTM1L-KO cells was not rescued by overexpression of CLPTM1(fig. S6, D to F). Therefore, CLPTM1 might have no functional overlap with CLPTM1L (*27*). To search for other GlcN-PI scramblases, we performed a further genome-wide CRISPR-Cas9 KO screen using CLPTM1L-KO HEK293 cells (fig. S7, A and B). However, reported ER scramblases, including TMEM16K, TMEM41B and VMP1 (*26, 28, 29*), were not identified (fig. S7C). These results suggest that yet unknown scramblases might contribute to GlcN-PI translocation in the absence of CLPTM1L.

**Fig. 4.**
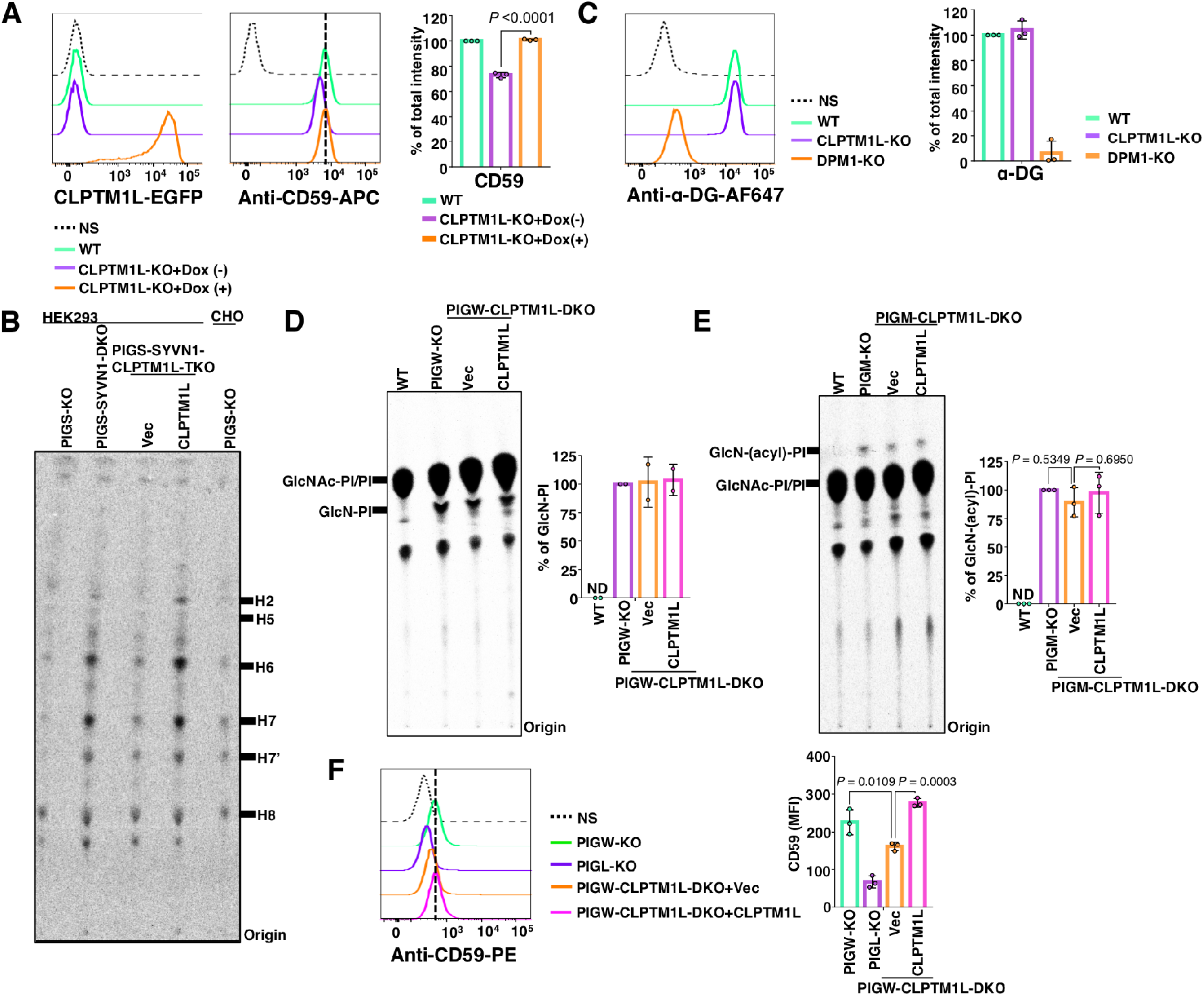
CLPTM1L is crucial for efficient utilization of GlcN-PI by the GPI biosynthetic pathway in the ER lumen. (**A**) Flow cytometry analysis of WT and CLPTM1L-KO HEK293 cells stably expressing CLPTM1L-mEGFP controlled by a doxycycline (Dox)-inducible Tet-On system (left) stained with anti-CD59 mAb (middle). CLPTM1L-mEGFP expression is induced by 1µg/mL of Dox. Right: Quantitative CD59 expression data from three independent experiments (mean ± SD, n = 3). (**B**) HPTLC analysis of GPI intermediates from cells metabolically labeled with [^3^H]Man for 1 hour. (**C**) Flow cytometry analysis of WT, DPM1-KO, and CLPTM1L-KO HEK293 cells stained with anti-α-dystroglycan (α-DG) mAb. DPM1 is essential for DPM biosynthesis. Right: Quantitative data of α-DG abundance from three independent experiments (mean ± SD, n = 3). (**D** and **E**) HPTLC analysis of GlcN-PI (D) or GlcN-(acyl)-PI (E) from cells metabolically labeled with [^3^H]inositol for 10 hours. Quantitative data from two (D) or three (F) independent experiments are shown on the right. (**F**) Flow cytometry analysis of PIGW-KO, PIGL-KO, and PIGW-CLPTM1L-DKO HEK293 cells stably expressing empty vector (Vec) or CLPTM1L stained with anti-CD59 mAb. Right: Quantitative data of MFI from three independent experiments (mean ± SD, n = 3). In (A), *P* value is from t test (unpaired and two-tailed). In (E) and (F), *P* values are from one-way ANOVA followed by Dunnett’s test for multiple comparisons.

To further characterize GPI biosynthesis in the absence of CLPTM1L, we analyzed Man-containing GPI intermediates after metabolic labeling of cells with [^3^H]Man. After cellular uptake, Man is converted to guanosine-diphosphate-Man, in turn to DPM and incorporated into GPI intermediates. For this experiment, we used PIGS-KO and PIGS-SYVN1-DKO HEK293 cells that are defective in GPI-transamidase required for attachment of preassembled GPIs to proteins. In these cells, synthesized GPIs remain un-linked to proteins, undergo structural maturation in the ER and Golgi similar to protein-linked GPI, and appear as free GPIs on the cell surface (*30*). We reported that GPI biosynthesis in PIGS-KO cells is enhanced several times after inactivation of the SYVN1-dependnt ER-associated protein degradation pathway (*31*), enabling GPI intermediates to be more easily determined after [^3^H]Man-labeling (Fig. 4B). To study the effects of CLPTM1L-defect on GPI biosynthetic intermediates, we generated PIGS-SYVN1-CLPTM1L-triple KO (TKO) HEK293 cells and those rescued by CLPTM1L cDNA (fig. S7D). KO of CLPTM1L caused partial decrease of all the detected Man-containing GPI intermediates (H2 to H6) and mature forms of GPIs (H7 and H8) that was restored by re-expression of CLPTM1L (Fig. 4B, and fig. S7D). No new and unusual GPI species were generated. Because the reduction of Man-labeled GPI intermediates could be caused by inefficient usage of DPM, and CLPTM1L might also mediate DPM scrambling considering its wide activity to various lipids in vitro (Fig. 3F and I), we addressed this possibility. Since DPM is required not only for GPI biosynthesis, but also for *N*-glycosylation, and protein O- and C-mannosylations, we assessed the status of α-dystroglycan, an O-mannosylation marker, and found that its level was not affected by KO of CLPTM1L in HEK293 cells (Fig. 4C, and fig. S7E), indicating that the reduction of Man-containing GPI intermediates was indeed caused by limited availability of GlcN-PI rather than DPM. These results support the conclusion that CLPTM1L-mediated GlcN-PI scrambling is a key step for the downstream stepwise reactions of GPI biosynthesis in the ER lumen.

We next analyzed GPI intermediates prior to mannosylation steps. Since GlcN-PI and GlcN-(acyl)- PI have very short lifetimes when the GPI biosynthetic pathway is ongoing, their detection by metabolic [^3^H]inositol labeling can be achieved in practice only by using cells defective in either inositol acylation (PIGW-KO cells) or the first mannosylation step (PIGM-KO cells). Therefore, we made PIGW-CLPTM1L-DKO cells and PIGM-CLPTM1L-DKO cells (fig. S7, F and G). The levels of GlcN-PI in PIGW-KO HEK293 cells was not appreciably affected by CLPTM1L KO (Fig. 4D), nor did we observe any significant decrease of GlcN-(acyl)-PI in PIGM-CLPTM1L-DKO cells compared to PIGM-KO cells (Fig.4E). Considering that detection of GlcN-(acyl)-PI by inositol labeling takes several hours of labeling, sufficient GlcN-PI is scrambled on this time frame even in the absence of CLPTM1L. The deficiency of one scramblase should only affect the rate at which GlcN-PI distributes between the two leaflets of the ER membrane, and therefore a significant change of the total amount of GlcN-PI or GlcN-(acyl)-PI is an unlikely effect of CLPTM1L-deficiency. Compared to wild-type cells, 5% of GPI-APs are expressed in PIGW-KO cells due to inefficient downstream reactions in the absence of inositol-acylation (*32*). Indeed, GlcN-PI scrambling mediated by CLPTM1L is important for GPI biosynthesis independent of the inositol acylation step, because the remaining CD59 expression in PIGW-KO cells made directly from GlcN-PI was still CLPTM1L-dependent (Fig. 4F).

Based on these results, we conclude that CLPTM1L is the major lipid scramblase mediating translocation of GlcN-PI across the ER membranes for GPI biosynthesis in mammalian cells. Although CLPTM1L is highly conserved in eukaryotes, it is notable that CLPTM1L homologs exist in only a few yeasts, but not *Saccharomyces*, suggesting the existence of other scramblases for GlcN-PI in *Saccharomyces* spp. (fig. S7H). This functional redundancy explains previous failures in identifying the enzyme for scrambling GlcN-PI. It is hypothesized that specific scramblases mediate the transmembrane translocation of other lipid glycoconjugates involved in post-translational modification of proteins in the ER lumen, and while the activity of these scramblases has been measured (*8*), their molecular identity has been elusive. Our discovery of CLPTM1L as a lipid scramblase for the GPI biosynthetic pathway validates this long-standing hypothesis, and provides insights into ER lumenal translocation of lipid glycoconjugates involved in protein glycosylation pathways.

## Supporting information

Table S1: Oligonucleotides used in this study.

Table S2: Guide RNA counts from CRISPR screen.

Table S3: Gene scores in unsorted versus sort2 PIGA-KO cells.

Table S4: Guide RNA counts from CRISPR screen.

Table S5: Gene scores in unsorted versus sort2 CLPTM1L-KO cells.

## Acknowledgments

We thank Dr. Tamao Endo and Dr. Hiroshi Manya (Tokyo Metropolitan Institute of Gerontology) for α-dystroglycan mAb (clone IIH6C4), and Dr. Toyoshi Fujimoto (Juntendo University) for expression plasmids of mouse *Ano10*. We thank Dr. Junji Takeda, Dr. Yusuke Maeda, and Dr. Yasu Morita (University of Massachusetts Amherst) for discussion, and Yuki Uchikawa and Yuko Kabumoto for assistance with cell sorting. We thank Keiko Kinoshita, Saori Umeshita, and Kae Imanishi for technical help. YW is supported by the IFReC Kishimoto Foundation Fellowship.

## Funding

Japan Society for the Promotion of Science (JSPS) KAKENHI Grant 21H02415 (TK)

Ministry of Education, Culture, Sports, Science and Technology (MEXT) of Japan KAKENHI Grant 17H06422 (TK)

A grant for Joint Research Project of the Research Institute for Microbial Diseases, Osaka University (MF and TK)

Max Planck Society (DVS and PHS)

RIKEN-Max Planck Joint Research Center for Systems Chemical Biology (PAG, DVS and PHS) Mizutani Foundation for Glycoscience (YMu)

## Author contributions

Conceptualization: YW, TK

Methodology: AKM, YMa, YMu, YI, PAG, DVS, PHS

Investigation: YW, AKM, YSL, MF, YMa

Visualization: YW

Funding acquisition: TK, MF, YMu, PAG, DVS, PHS

Project administration: YW, TK

Supervision: TK, AKM

Writing – original draft: YW, TK

Writing – review & editing: YW, YSL, DVS, PHS, YI, MF, YMa, YMu, AKM, TK

## Data and materials availability

All data are available in the main text or the supplementary materials.

**Fig. S1.**
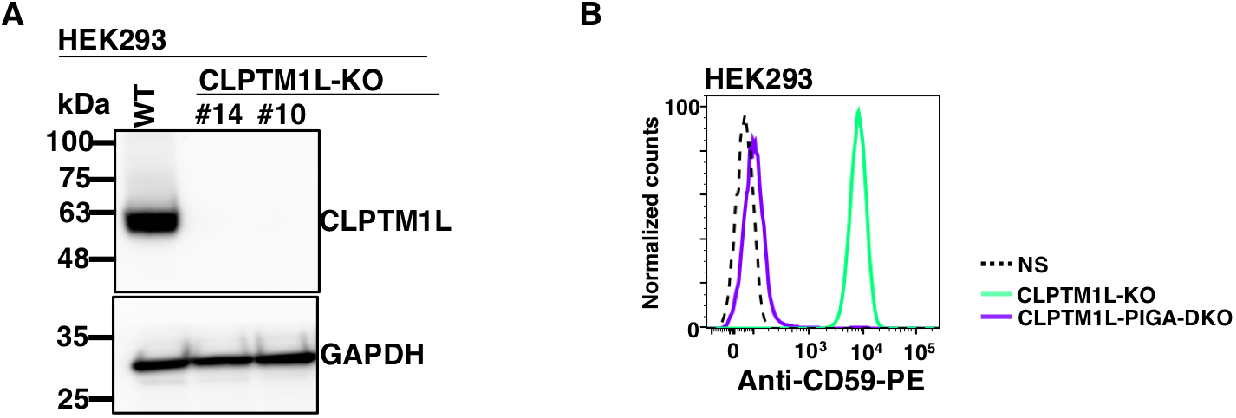
Data supporting Figures 1 and 2. **(A)** Western blotting of endogenous CLPTM1L. Lysates of wild-type (WT) and two clones of CLPTM1L-KO HEK293 cells were analyzed by western blotting. GAPDH, a loading control. **(B)** Confirmation of PIGA-KO in CLPTM1L-KO HEK293 cells (clone #10) to generate CLPTM1L-PIGA-DKO cells. Cells stained with PE-labeled anti-CD59 mAb were analyzed by flow cytometry.

**Fig. S2.**
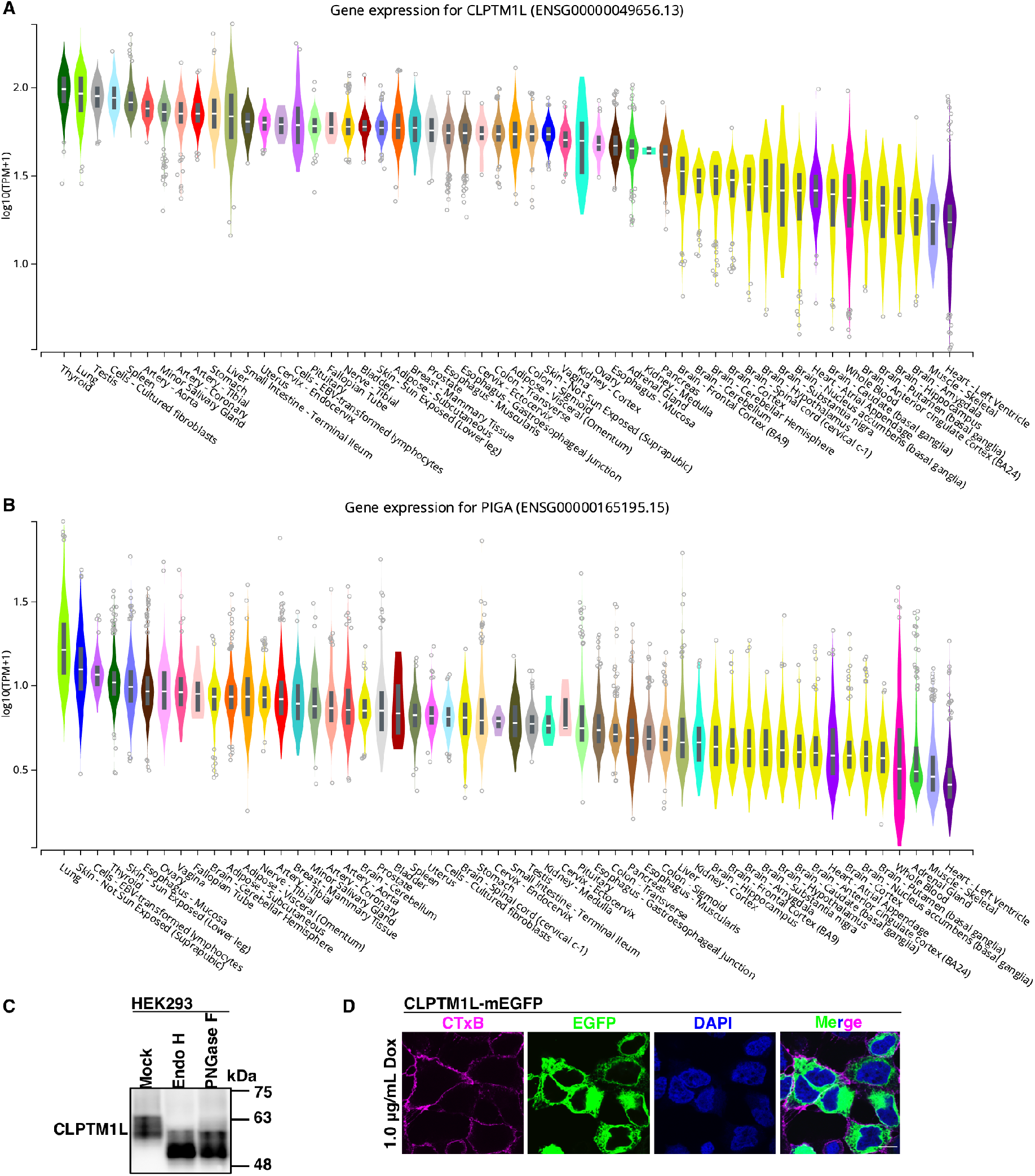
Data supporting Figures 1 and 2. **(A and B)** Gene expression analysis for CLPTM1L (A) and PIGA (B) across different tissues, taken from GTEx Portal (*33*). **(C)** Western blotting of endogenous CLPTM1L. Lysates of HEK293 cells treated with or without EndoH or PNGase F were analyzed by western blotting. **(D)** Representative images of CLPTM1L-KO HEK293 cells expressing CLPTM1L-mEGFP induced by doxycycline (Dox). Cells were labeled with CTxB to detect a plasma membrane marker, GM1a ganglioside. Scale bar, 10 μm.

**Fig. S3.**
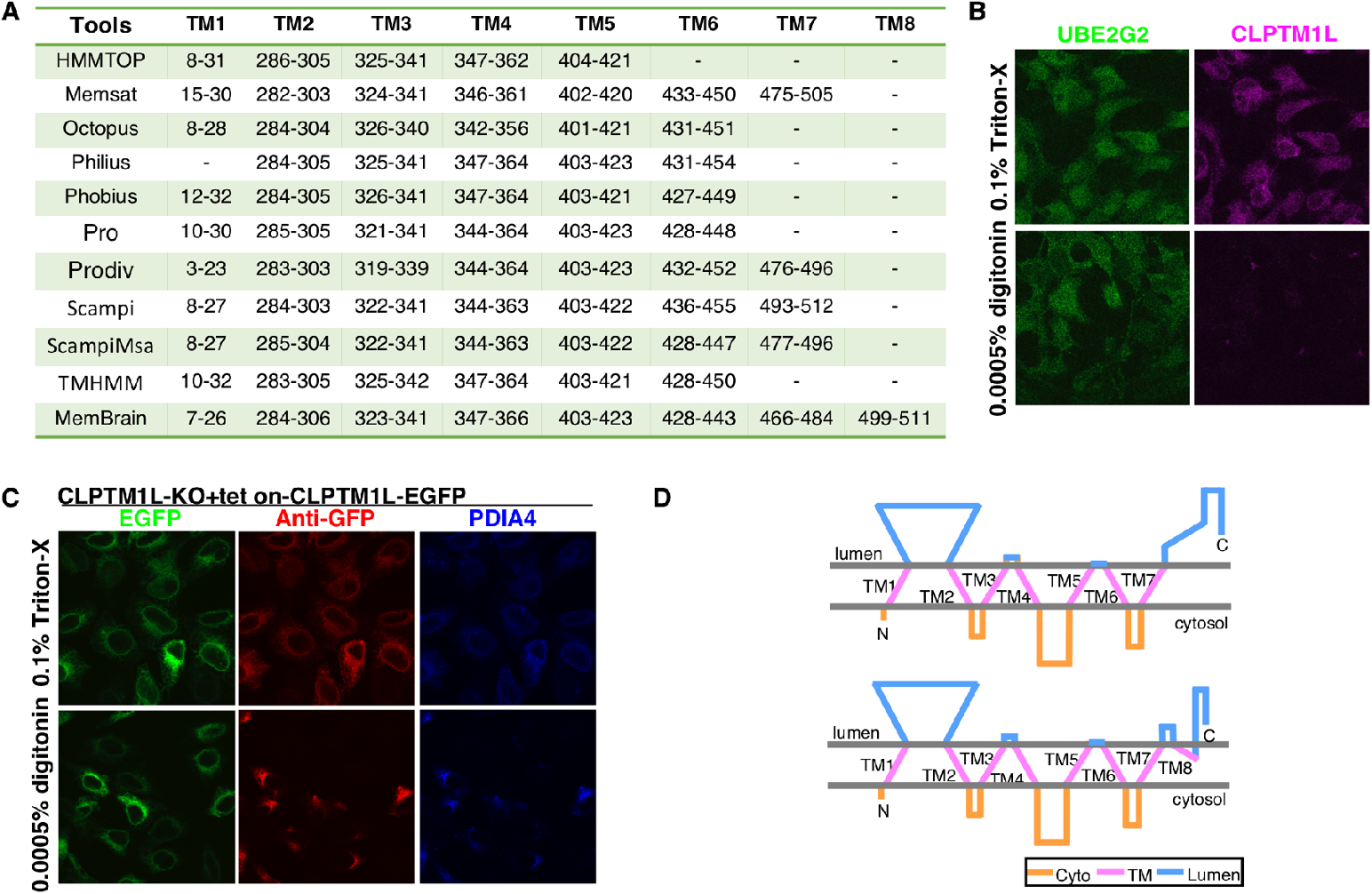
Data supporting Figure 2. **(A)** Transmembrane domain prediction using various algorithms (*34*–*36*). **(B)** Representative images of HEK293 cells permeabilized by 0.0005% digitonin or 0.1% Triton X-100, and endogenous CLPTM1L was detected using antibody specifically recognizing the region between TM1 and TM2 of CLPTM1L. UBE2G2, a cytosolic protein, was detected by anti-UBE2G2 mAb. **(C)** Representative images of CLPTM1L-KO HEK293 cells expressing CLPTM1L-mEGFP induced by Dox, permeabilized with 0.0005% digitonin or 0.1% Triton X-100. CLPTM1L-mEGFP was visualized by green fluorescence and the C-terminal mEGFP was detected using anti-GFP antibody. PDIA4, an ER lumenal protein, was detected by anti-PDIA4 mAb. **(D)** Models of the predicted topology of CLPTM1L with 7 transmembrane (TM) domains (upper) or 8 TM domains (lower) in the ER membrane. TM domains are indicated by numbers. Cyto, cytoplasm.

**Fig. S4.**
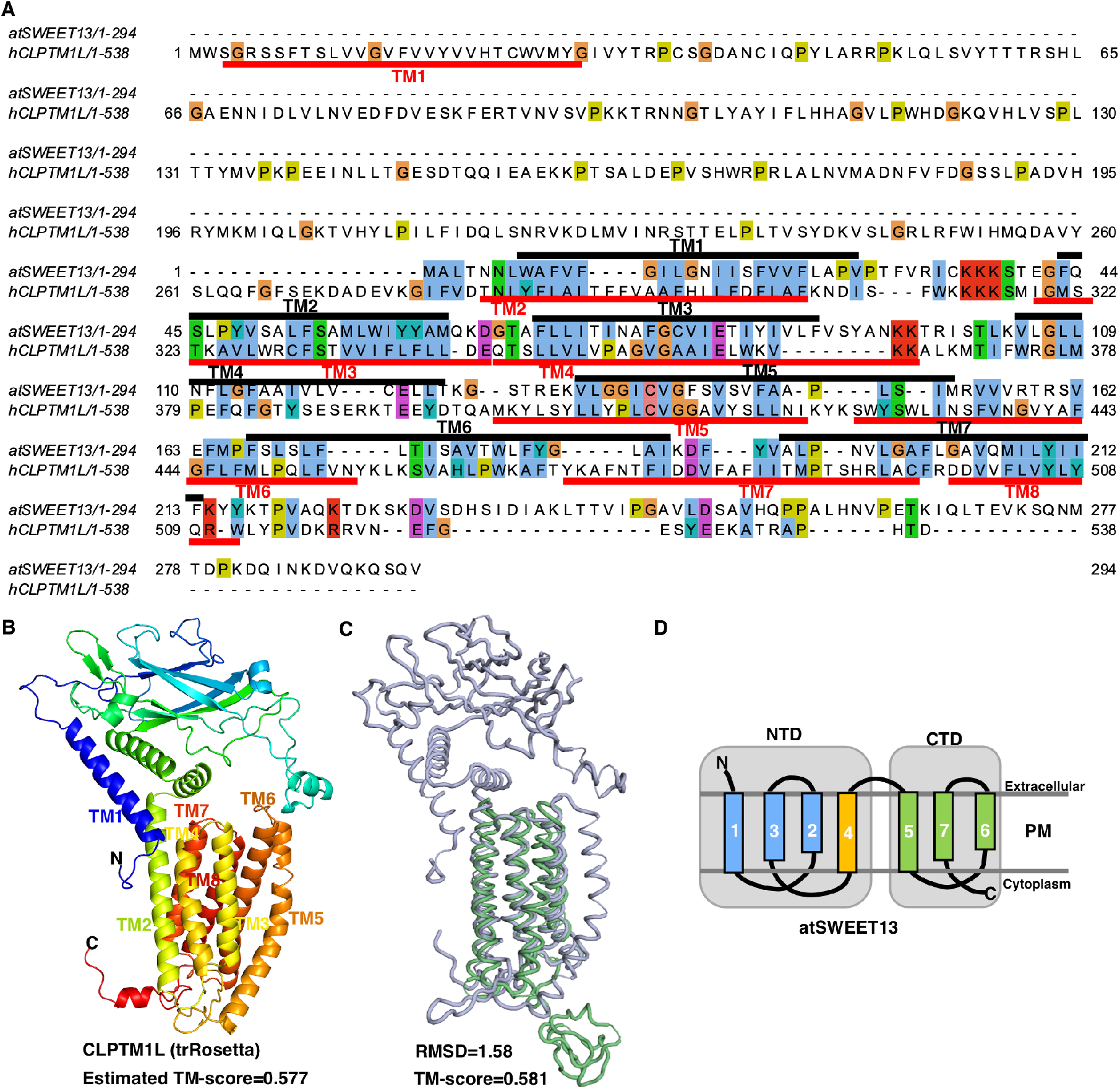
Data supporting Figure 2. **(A)** Sequence alignment of human CLPTM1L and *Arabidopsis thaliana* SWEET13 (atSWEET13). The TM domains are underlined in red for CLPTM1L or overlined in black for atSWEET13. Sequences are from Uniprot; human CLPTM1L (Q96KA5); and atSWEET13 (Q9FGQ2). The sequence alignment was generated using Clustal Omega program and colored by the Clustal style in the Jalview (*37, 38*). **(B)** Three-dimensional structural model of human CLPTM1L predicted by trRosetta. **(C)** Structural comparison of predicted CLPTM1L structure and atSWEET13. CLPTM1L is shown in light blue, and atSWEET13 (PDB entry 5XPD) in green. **(D)** Topology of atSWEET13 in the plasma membrane (PM). AtSWEET13 has 7 TM domains in a 3+1+3 repeat arrangement in the PM. NTD, N-terminal domain; CTD, C-terminal domain. The TM domains are indicated by numbers in (A), (B) and (D).

**Fig. S5.**
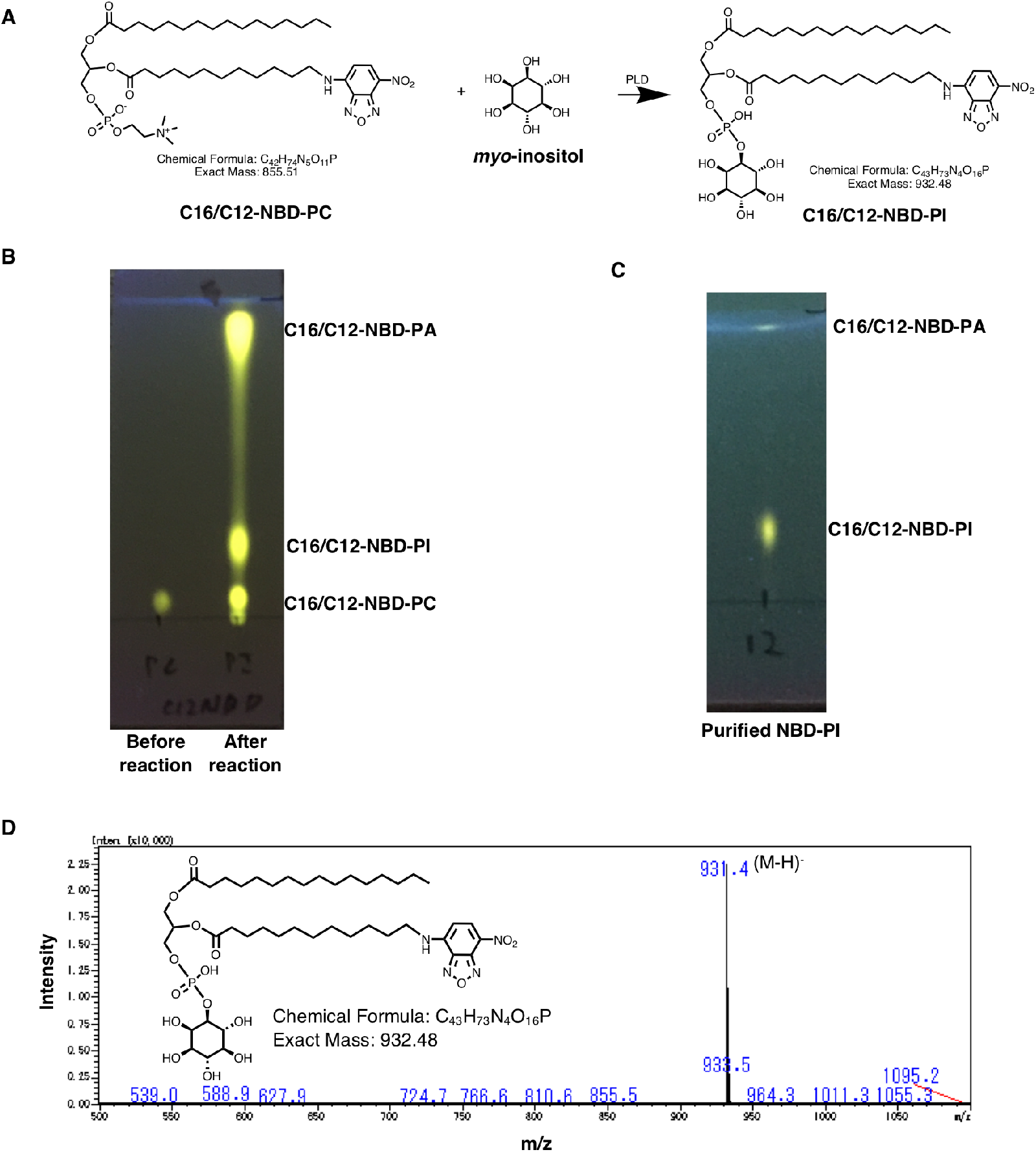
Data supporting Figure 3. **(A)** Reaction scheme of PLD mediated transphosphatidylation of C16/C12-NBD-PC to generate C16/C12-NBD-PI. **(B and C)** TLC analysis of lipids after enzyme reaction (B) and after purification (C). Solvent, chloroform/ petroleum ether/ methanol/ acetic acid (4:3:2:1, by volume), was used to develop the TLC plate, and the spots were visualized under UV light at 365 nm. **(D)** Liquid chromatography–mass spectrometry analysis of the purified C16/C12-NBD-PI. Mass spectrum of the chromatogram peak showed [M-H]-ions at m/z 931.4, confirming the presence of C16/C12-NBD-PI.

**Fig. S6.**
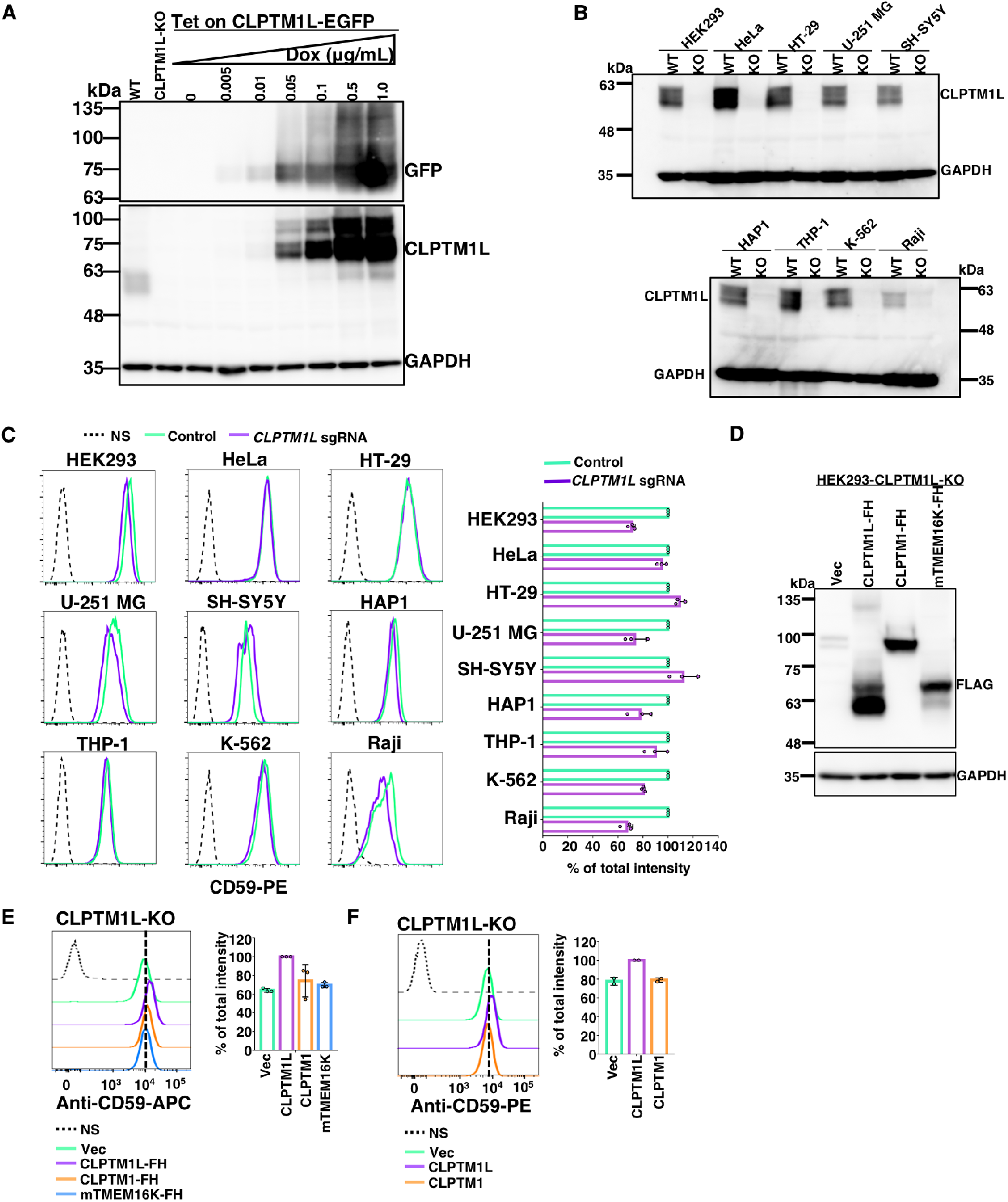
Data supporting Figure 4. **(A)** Western blotting of CLPTM1L-mEGFP induced by Dox at different concentrations. CLPTM1L-mEGFP was detected by anti-GFP and anti-CLPTM1L antibodies. **(B)** Western blotting of endogenous CLPTM1L of various human cell lines. Lysates of wild-type (WT) and CLPTM1L-deficient (KO) cells were analyzed by western blotting. **(C)** Flow cytometry analysis of various human cultured cell lines with CLPTM1L-deficeincy. Cells stably expressing lentiCRISPR v2 with CLPTM1L-targeting sgRNAs were stained with anti-CD59 mAb. **(D)** Western blotting of CLPTM1L-FLAG-6His (FH), CLPTM1-FH and mouse TMEM16K-FH induced by Dox in CLPTM1L-KO HEK293 cells. GAPDH, a loading control in (A), (B), and (D). **(E and F)** Flow cytometry analysis of CLPTM1L-KO HEK293 cells stained with anti-CD59 mAb. In (E), overexpression of CLPTM1L-FH, CLPTM1-FH, and mTMEM16K-FH controlled by Tet-On system was induced by 1 µg/mL Dox. In (F), retroviral vector-mediated overexpression of CLPTM1L or CLPTM1 in CLPTM1L-KO HEK293 cells. Quantitative data from three (C and E) or two (F) independent experiments are shown on the right.

**Fig. S7.**
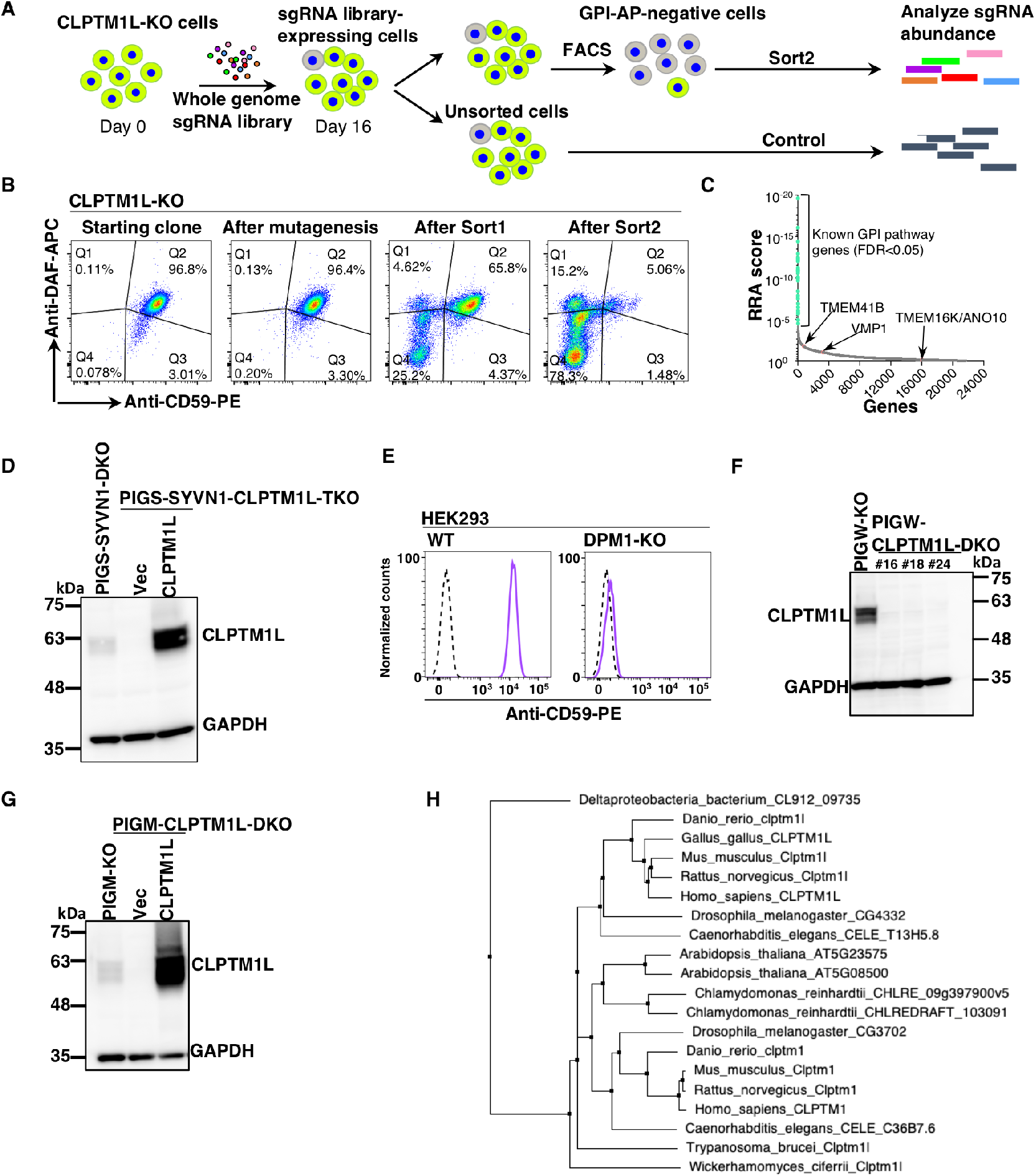
Data supporting Figure 4. **(A)** Scheme depicting a FACS-based genome-wide CRISPR screen for genes involved in GPI biosynthesis using CLPTM1L-KO HEK293 cells. **(B)** Flow cytometry analysis of CLPTM1L-KO HEK293 cells during the FACS-based genome-wide CRISPR screen. Starting clone, cells after mutagenesis, sort1 cells, and sort2 cells were stained with anti-CD59 and anti-CD55 antibodies. **(C)** Gene scores in unsorted versus sorted CLPTM1L-KO cells. Known GPI biosynthetic pathway genes are shown in green. Reported ER scramblase genes including TMEM41B, VMP1, and TMEM16K/ANO10 were shown in pink. **(D)** Western blotting of CLPTM1. Lysates of PIGS-SYVN1-DKO and PIGS-SYVN1-CLPTM1L-TKO HEK293 cells stably expressing empty vector (Vec) or CLPTM1L were analyzed by western blotting. **(E)** Confirmation of DPM1-KO in HEK293 cells. WT and DPM1-KO HEK293 cells stained with PE-labeled anti-CD59 mAb were analyzed by flow cytometry. **(F)** Confirmation of CLPTM1L-KO in PIGW-KO cells by western blotting. PIGW-CLPTM1L-DKO (clone #18) was used for the downstream analyses. **(G)** Lysates of PIGM-KO and PIGM-CLPTM1L-DKO HEK293 cells stably expressing empty vector (Vec) or CLPTM1L were analyzed by western blotting. GAPDH, a loading control in (D), (F), and (G). **(H)** Phylogenetic tree of CLPTM1 family members.

**Table S1**.

Oligonucleotides used in this study.

**Table S2**.

Guide RNA counts from CRISPR screen described in Fig. 1 and their related text.

**Table S3**.

Gene scores in unsorted versus sort2 cells described in Fig. 1 and their related text.

**Table S4**.

Guide RNA counts from CRISPR screen described in Fig. S7 and their related text.

**Table S5**.

Gene scores in unsorted versus sort2 cells described in Fig. S7 and their related text.

## Materials and Methods

### Antibodies and chemicals

Anti-CD55 (clone IA10, BD Biosciences) and anti-CD59 (clone 5HA) mAbs were described previously (*39*). Commercial primary antibodies (catalog number, commercial source) used in this study were: GAPDH (MA1-22670, Thermo Fisher Scientific), CLPTM1L (HPA014791, Atlas Antibodies), UBE2J1 (sc-377002, Santa Cruz Biotechnology), UBE2G2 (sc-393780, Santa Cruz Biotechnology), DYKDDDDK/FLAG (014-22383, Wako), GFP (ab5450, Abcam), GFP (ab290, Abcam), PDIA4 (5033S, Cell Signaling Technology), α-dystroglycan (05-593, Merck Millipore), CD59 (304708, BioLegend).

GlcNAc-PI and GlcN-PI used in this study were chemically synthesized as recently described (Paula A. Guerrero et al., manuscript under review), the PI portion is 1-stearoyl-2-oleoyl-sn-glycero-3-phospho-1D-myo-inositol. Commercial chemicals (catalog number, commercial source) used in this study were: doxycycline (631311, TaKaRa), tunicamycin (202-08241, Wako), DTT (P2325, Thermo Fisher Scientific), digitonin (D5628, Sigma), DSP (22585, Thermo Fisher Scientific), n-dodecyl β-D-maltoside (DDM) (14239-41, Nacalai Tesque), ATP disodium salt (A2383, Sigma-Aldrich), GTP (G8877, Sigma-Aldrich), bovine brain phosphatidylserine (P7769, Sigma-Aldrich), egg phosphatidylcholine (840051C, Avanti Polar Lipids), 18:1 PS/DOPS (840035P, Avanti Polar Lipids), 16:0-12:0 NBD PC (810131P, Avanti Polar Lipids), 18:1 NBD PE (810145P, Avanti Polar Lipids), sodium dithionite (S0562, TCI), D-[2-^3^H]mannose (ART0120A, American Radiolabeled Chemicals), myo-[2-^3^H]inositol (NET114A001MC, PerkinElmer), UDP-[6-^3^H]GlcNAc (ART 1136, American Radiolabeled Chemicals), phosphatidylinositol, L-α-[myo-inositol-2-^3^H(N)] (ART 0184, American Radiolabeled Chemicals).

### Cells and culture

HEK293 cells (ATCC CRL-1573), HT-29 cells (ATCC HTB-38), U-251 MG cells, SH-SY5Y cells (ATCC CRL-2266), HeLa cells (ATCC CCL-2), and their knockout (KO) derivatives were cultured in DMEM with high glucose (Nacalai Tesque) containing 10% heat-inactivated fetal bovine serum (FBS) (172012-500ML, Sigma). Raji cells (ATCC CCL-86), THP-1 cells (ATCC TIB-202), K-562 cells (ATCC CCL-243), and their derivatives were cultured in RPMI 1640 medium (Nacalai Tesque) with 10% heat-inactivated FBS. HAP1 cells (kindly provided by Thijn Brummelkamp) and their KO derivatives were cultured in IMDM medium (Nacalai Tesque) supplemented with 10% heat-inactivated FBS. CHO K1 cells (ATCC CCL-61) and their derivatives were maintained in DMEM/F-12 (Nacalai Tesque) supplemented with 10% heat-inactivated FBS. All cell lines were maintained in an incubator at 37 °C and 5% CO_2_.

### FACS-based genome-wide CRISPR screen

#### CRISPR screens were performed as recently described (*31*)

Lenti-X 293T (Clontech) cells were seeded at 70% confluence to one 10 cm-dish and incubated for 10 hours to reach about 90% confluence before transfection. To produce lentivirus, the human GeCKOv2 pooled plasmids (lentiCRISPRv2) were co-transfected with lentiviral packaging plasmids pLP1, pLP2, and pLP/VSVG (Thermo Fisher) into Lenti-X 293T cells. For one 10 cm-dish, plasmids including pLP1 (6.3 μg), pLP2 (2.1 μg), pLP/VSVG (3.2 μg) and lentiCRISPRv2 (4.2 μg library A and 4.2 μg library B) were diluted in 1.25 mL of OptiMEM (Thermo Fisher) to prepare DNA reagent. Forty microliters of 2 mg/mL PEI “Max” (Polyscience) was diluted in 1.25 mL of OptiMEM and incubated for 5 min at room temperature (RT). This PEI “Max” reagent was added to the DNA reagent with vortex. The mixture was incubated at RT for 30 min and added to the cells. After 12 hours, the media was changed to 10 mL of pre-warmed DMEM supplemented with 10% FBS. About 24 hours after the media change, the viral media was collected and 10 mL of pre-warmed DMEM was added to the cells. The supernatant was filtered through a Millex-HV syringe filter (SLHV033R, Merck Millipore). About 48 and 72 hours after the media change, the viral media was collected and filtered. Finally, 30 mL of viral media was combined and stored at 4 °C for functional titration and large-scale CRISPR screens.

For functional titration, about 2.5 × 10^5^ CLPTM1L-KO or PIGA-KO HEK293 cells per well were plated in 6-well plates (IWAKI) and maintained for 36 hours before adding the viral media. Different volumes of viral supernatant (50, 100, 150, 200, 300, 400, and 500 μL) were added to each well along with a no virus control with 7 μg/mL polybrene (Sigma-Aldrich). Cells were incubated overnight, and media were changed to pre-warmed DMEM. At 24 hours post-transduction, cells in all wells were split to a 1:5 ratio to prevent any well from confluence, and 10, 20, 25, 30, and 50% confluent controls were treated similarly. At 2 days and a half post-transduction, except for the confluent controls, cells were maintained in DMEM supplemented with 0.5 μg/mL puromycin (Sigma Aldrich). Cells were split 1:5 to prevent any well from confluence. At 10 days post-transduction, all cells in the no virus control with puromycin were floating. The viral volume that results in 30% cell survival (MOI ≈ 0.3) in 0.5 μg/mL puromycin was determined by comparing with the 30% confluence control.

For large-scale CRISPR screen using PIGA-KO HEK293 cells as the parent cells, 6 × 10^6^ cells were plated in one 15 cm-dish 1 day before adding the virus, 12 × 15 cm-dishes were prepared. Around 1.2 × 10^7^ cells per dish were transduced with 7 mL of viral supernatant. At 2 days and a half post-transduction, the infected cells were maintained with 1 μg/mL puromycin. To keep the complexity of gRNA library, cells were combined, and a minimum of 6 × 10^7^ cells were placed for culture. At about 2 weeks post-transduction, about 8 × 10^7^ cells were maintained in the DMEM without FBS containing 1.5 µM GlcNAc-PI (7.5 mM compound stock in DMSO) overnight. About 1.2 × 10^8^ cells were then harvested and stained by anti-CD59 mAb (clone 5H8), followed by staining with PE-labeled goat anti-mouse IgG antibody (405307, BioLegend). For large-scale CRISPR screen using CLPTM1L-KO HEK293 cells as the parent cells, 3.5 × 10^6^ cells were plated in one 15 cm-dish 36 hours before adding the virus. 7 × 15 cm-dishes in total were prepared. Around 1 × 10^7^ cells per dish were transduced with 6 mL of viral supernatant. At 2 days and a half post-transduction, the infected cells were maintained with 0.5 μg/mL puromycin until the cells were expanded to 2.4 × 10^8^. At about 2 weeks post-transduction, about 1 × 10^8^ cells were harvested and stained by PE-labeled anti-CD59 mAb. After washing by cold PBS, the cells were resuspended in Hanks’ Balanced Salt Solution (H6648, Sigma-Aldrich), and sorted by FACSAria (BD) to collect CD59 mAb staining-negative cells. On the same day, pellet of at least 6 × 10^7^ cells without sorting was stocked at –80 °C. After the first sorting, the cells were maintained in DMEM supplemented with 0.25 μg/mL puromycin. For the second sorting, at least 1 × 10^7^ cells were prepared. The sort2 cells were maintained in the medium until expanded to 2 × 10^7^. Pellets of 2 × 10^7^ sort2 cells were stored at –80 °C for genomic DNA purification.

### Analysis of CRISPR screen

Approximately 6 × 10^7^ control cells and 2 × 10^7^ sort2 cells were used for genomic DNA extraction by Wizard Genomic DNA Purification Kit (A1125, Promega). Approximately 325 μg of genomic DNA from control cells and 60 μg of genomic DNA from sort2 cells were used for amplification of gRNA. PCR (27 cycles) was performed to amplify the sgRNAs using Ex Taq Polymerase (RR001A, TaKaRa), in total 65 tubes for control cells and 12 tubes for sort2 cells (Oligos for amplification of sgRNAs are in Table S1). All the PCR products were combined and mixed well, and 675 μL PCR products for control cells and 320 μL were applied to the 2% agarose gel for purification using QIAquick Gel Extraction Kit (QIAGEN). The PCR products of control and sort2 were concentrated by using Amicon Ultra-0.5 10K (UFC501096, Merck Millipore), and analyzed by pair-end sequencing using NovaSeq 6000 system (Illumina). Raw data were processed for sgRNA counting using the Python scripts. The sequencing reads were demultiplexed using the 5 bp adapter by cutadapt version 2.8 (*40*). The adapters of the demultiplexed reads were then trimmed by bowtie2 (*41*), obtaining 20 bp gRNA sequences. These sgRNA sequences were mapped to the sequences of Human GeCKO v2 sgRNA library, and the total number of sgRNA counts were obtained by using the MAGeCK workflow version 0.5.8, the robust rank aggregation (RRA) values and p-values were determined using the MAGeCK algorithm (*42*). The MAGeCK outputs of the two CRISPR screens described in this study were shown in Table S2 to S5.

### Recombinant DNA

Human GeCKOv2 CRISPR knockout pooled library (Pooled Library #1000000048),pX330-U6-Chimeric_BB-CBh-hSpCas9 (Addgene plasmid # 42230), and lentiCRISPR v2 (Addgene plasmid # 52961) were gifts from Feng Zhang (*9, 43*). pX330-mEGFP plasmid was generated from pX330-U6-Chimeric_BB-CBh-hSpCas9 (*44*). The plasmids of the PiggyBac Tet-On expression system including pPB-tetIRB-tetON-BSD and pPB-tetON-BSD-tightFF were gifts from Yusuke Maeda (Osaka University). The hyperactive PB transposase expression vector (pCMV-hyPBase) was a gift from Kosuke Yusa (*45*). Mouse *Ano10/Tmem16k* cDNA was a gift from Toyoshi Fujimoto (*46*). Retroviral vector pLIB2-BSD-hCLPTM1 was previously described (*47*). Human CLPTM1L cDNA was amplified by PCR from a human cDNA library and cloned into the vector pLIB2-BSD to construct pLIB2-BSD-hCLPTM1L. Plasmid pLIB2-BSD-hCLPTM1L-FLAG-6His was constructed by sub-cloning the EcoRI-MluI fragment of hCLPTM1L from pLIB2-BSD-hCLPTM1L into the same sites of pLIB2-BSD-hB3GALT4-FLAG-6His (*31*). To construct pLIB2-Hyg-hCLPTM1L-3HA, EcoRI-MluI fragment of hCLPTM1L was cloned into pME-3HA, and the EcoRI-MluI fragment of hCLPTM1L-3HA was cloned into pLIB2-Hyg. To construct pPB-tetON-hCLPTM1L-FLAG-6His-BSD-tightFF plasmid, hCLPTM1L-FLAG-6His was amplified by PCR and cloned into pPB-tetON-BSD-tightFF plasmid by In-fusion cloning (Clontech). To construct pPB-tetON-Tmem16k-FLAG-6His-BSD-tightFF plasmid, mouse *Ano10/Tmem16k* cDNA was amplified by PCR from pMXs-IRES-Puro-mTMEM16K-FLAG, then cloned into pPB-tetON-hCLPTM1L-FLAG-6His-BSD-tightFF plasmid by In-fusion cloning. Other plasmids including pPB-tetON-hCLPTM1-FLAG-6His-BSD-tightFF and pPB-tetIRB-tetON-CLPTM1L-mEGFP-BSD were also constructed by In-fusion cloning. Oligos for generation of the plasmids used in this study were shown in Tables S1.

### Gene knockout

Gene knockout was achieved using the CRISPR/Cas9 system. To generate HEK293 knockout clones, sgRNAs targeting to the exon regions of each gene was cloned into pX330-mEGFP plasmids. CLPTM1L-KO HEK293 clones including CLPTM1L-KO, PIGS-SYVN1-CLPTM1L-TKO, PIGW-CLPTM1L-DKO, and PIGM-CLPTM1L-DKO were validated by PCR to confirm a large deletion of the amplicon from CLPTM1L gene and immunoblotting to confirm the loss of endogenous CLPTM1L protein. To generate CLPTM1L-PIGA-DKO cells, PIGA-KO from CLPTM1L-KO cells were confirmed by FACS analysis to confirm complete loss of CD59 expression. DPM1-KO HEK293 cells were confirmed by FACS analysis of cell surface CD59 and α-dystroglycan expression. To knockout CLPTM1L in various human cell lines, sgRNAs targeting to the exon regions of CLPTM1L were cloned into lentiCRISPR v2 vector. Cells were transduced with lentiCRISPR v2 lentivirus and maintained in their medium containing 1-2 μg/mL puromycin. The polyclonal cell populations with CLPTM1L-deficiency were validated by western blotting to confirm the loss of CLPTM1L protein in each cell line. Other KO HEK293 cells including PIGA-KO, PIGW-KO, PIGL-KO, and PIGM-KO cells were recently described (Paula A. Guerrero et al., manuscript under review). sgRNA sequences used in this study were in Tables S1.

### Western blotting

Cell pellets were washed twice with ice-cold PBS, and lysed in RIPA lysis buffer (9806S, Cell Signaling Technology) supplemented with cOmplete protease inhibitors (Roche) on ice for 30 min, followed by centrifugation at 17,900 × *g* for 15 min at 4 °C. Supernatants were mixed with 4 × SDS-sample buffer containing 5% β-mercaptoethanol and incubated on ice for 30 min. In some cases, the denatured samples were incubated at 37 °C with or without Endo Hf (P0703S, New England Biolabs) or PNGase F (P0704S, New England Biolabs) for 2 hours prior to electrophoresis. Samples were then resolved on SDS-PAGE gels, then transferred to PVDF membranes (Merck Millipore) for immunoblotting. The membranes were blocked at RT in Tris-buffered saline containing 5% nonfat milk and 0.1% Tween-20 (TBS-T) for 1 hour, followed by incubation with primary antibodies in Can Get Signal Solution 1 (TOYOBO) at RT for 1 hour. After washing three times with TBS-T buffer, the membranes were then incubated with horseradish peroxidase labeled secondary antibodies in Can Get Signal Solution 2 (TOYOBO) for 1 hour at RT. After washing with TBS-T, blots were exposed to Amersham ECL Prime Western Blotting Detection Reagent (GE Healthcare), and images were captured using ImageQuant LAS 4000 Mini (GE Healthcare).

### Retrovirus-mediated gene overexpression

PLAT-GP retroviral packaging cells were seeded at about 60% confluency on 6-well plate overnight before transfection to reach ∼90% confluence. Cells were transfected with pLIB2-BSD plasmid bearing cDNA of human CLPTM1L or CLPTM1 by using PEI “Max” transfection reagent (Polyscience) and cultured for 12 hours. Cells were incubated in medium containing 10 mM sodium butyrate (Sigma Aldrich) overnight. Cells were incubated in fresh medium at 32 °C for 1 day. Viral medium was added to cells, followed by centrifugation at 1,100 × g at 32 °C for 4 hours. These cells were cultured at 32 °C overnight. On the next day, culture medium was replaced with fresh medium, and the cells were cultured at 37 °C for another 2 days. Cells were maintained in the medium with 10 μg/mL blasticidin (ant-bl-1, InvivoGen) for at least two weeks to achieve complete antibiotic selection before using for other experiments.

### Tet-On tetracycline-inducible gene overexpression

Expression of CLPTM1L-mEGFP, CLPTM1L-FLAG-6His, CLPTM1-FLAG-6His, or mouse Ano10/ Tmem16k-FLAG-6His in HEK293 cells or Expi293F cells was achieved through the PiggyBac Tet-On expression system. Cells were transfected with pCMV-hyPBase, and pPB-tetIRB-tetON-BSD plasmid bearing CLPTM1L-mEGFP or pPB-tetON-BSD-tightFF plasmid bearing CLPTM1L-FLAG-6His using X-tremeGENE 9 DNA transfection reagent (Roche), and incubated at 37 °C for 2 days. Cells were then maintained in the medium containing 10 μg/mL blasticidin for two weeks, achieving complete antibiotic selection before using for other experiments.

### Flow cytometry

Cells stained with biotin-labeled anti-CD59 mAb (clone 5H8), biotin-labeled anti-CD55/DAF mAb (clone IA10), or anti-α-dystroglycan (clone IIH6C4) in FACS buffer (PBS containing 1% BSA and 0.1% NaN_3_) were incubated on ice for 25 min. Cells were then washed twice in FACS buffer followed by staining with Alexa Fluor 647-conjugated goat anti-mouse IgM (ab150123, Abcam) for α-dystroglycan mAb, allophycocyanin (APC)-labeled streptavidin (405207, BioLegend) for biotin-labeled anti-CD59 or anti-CD55 mAb, or phycoerythrin (PE)-labeled anti-CD59 mAb (304708, BioLegend) in FACS buffer. After twice washing by FACS buffer, cells were analyzed by using the BD FACSCanto II.

### Immunofluorescence

For cell surface staining by CTxB, CLPTM1L-KO HEK293 cells expressing Tet-On system controlled CLPTM1L-mEGFP were allowed to adhere to gelatin-coated coverslips overnight at 37 °C. Expression of CLPTM1L-mEGFP was induced by 1 µg/mL doxycycline (TaKaRa) and maintained in DMEM supplemented with Tet System Approved FBS (631107, TaKaRa) for 2 days. Cells were rinsed with ice-cold PBS three times and incubated with 1 μg/mL Alexa Fluor 594-conjugated CTxB (C34777, Thermo Fisher Scientific) on ice for 20 min. After washing with ice-cold PBS, the cells were immediately fixed with 4% paraformaldehyde in PBS for 20 min at RT and incubated with 50 mM ammonium chloride for 10 min. Coverslips were then washed with PBS three times and mounted in ProLong Gold antifade reagent with DAPI (P36941, Thermo Fisher Scientific). Images were taken on an Olympus FV1000 laser scanning confocal microscope with a UPLSAPO oil lens (×100 magnification and 1.4 NA).

To detect endogenous CLPTM1L in HEK293 cells, wild-type and CLPTM1L-KO HEK293 cells were seeded on the coverslips, fixed with 4% paraformaldehyde and permeabilized in blocking buffer with 0.1% saponin for 1 hour at RT, and then incubated with 1 μg/mL rabbit anti-CLPTM1L (1:500), and mouse anti-UBE2J1 (1:250) diluted in blocking solution with 0.1% saponin and 5% goat serum (Gibco) for 1 hour. After three washes in PBS for 30 min, coverslips were incubated with Alexa Fluor 594-conjugated goat anti-rabbit IgG (1:500) and Alexa Fluor 488-conjugated goat anti-mouse IgG (1:500) for 1 hour. Images were taken on an Olympus FV1000 laser scanning confocal microscope with a UPLSAPO oil lens (×100 magnification and 1.4 NA).

To analyze the orientation of endogenous CLPTM1L, HEK293 cells were permeabilized in blocking solution with 0.0005% digitonin for 5 min at RT, and blocked in FACS buffer containing 5% normal donkey serum (Jackson ImmunoResearch) with or without 0.1% Triton X-100 for 30 min. Cells were stained with rabbit anti-CLPTM1L (1:500) and mouse anti-UBE2G2 (1:200) mAbs in FACS buffer for 2 hours at RT, followed by staining with Cy3-conjugated donkey anti-mouse IgG (1:500) and Alexa Fluor 647-conjugated donkey anti-rabbit IgG (1:500) in FACS buffer for 1 hour at RT. To analyze CLPTM1L-mEGFP induced by 1 µg/mL doxycycline, cells stained with goat anti-GFP and rabbit anti-PDIA4 (1:500) antibodies were incubated with Alexa 594-conjugated donkey anti-goat IgG (1:500) and Alexa 647-conjugated donkey anti-rabbit IgG (1:500) in FACS buffer with 2.5% normal donkey serum. After washing, coverslips were mounted in ProLong Diamond anti-fade reagent (P36970, Thermo Fisher Scientific). Images were taken on an Olympus FV1000 laser scanning confocal microscope with a UPLSAPO oil lens (×60 magnification and 1.35 NA).

### Immunoprecipitation

Immunoprecipitation was described previously (*47*). Briefly, CLPTM1L-KO HEK293 cells expressing CLPTM1L-FLAG-6His (CLPTM1L-FH) and CLPTM1L-3HA or empty vector were cultured in 10-cm dishes, harvested and washed in cold PBS. Cells were lysed in 600-μL of NP-40 lysis buffer (25 mM HEPES, pH 7.4, 150 mM NaCl, 1% NP-40, cOmplete protease inhibitor, and 1 mM PMSF) with rotation at 4 °C for 1 hour. After centrifugation at 12,000 × *g* for 15 min at 4 °C, 30 μL of each supernatant was kept as total lysate and the rest of each supernatant was transferred to new tubes containing 20 μL of anti-Flag M2 beads (A2220-25ML, Sigma Aldrich) prewashed in lysis buffer. After rotation at 4 °C for 2 hours, the beads were washed four times with lysis buffer. The proteins were then eluted from the beads with 100 μL of lysis buffer containing 100 μg/mL 3 × FLAG peptide (F4799, Sigma Aldrich), mixed with sample buffer and incubated on ice for 30 min. Samples were then subjected to SDS-PAGE and analyzed by western blotting.

### Cross-linking

CLPTM1L-KO HEK293 cells expressing CLPTM1L-FH were incubated in ice-cold PBS containing various concentrations of DSP (0-2mM) for 30 min. To quench the reaction, cells were incubated in ice-cold PBS containing 100 mM Tris-HCl for 20 min. Cells were washed twice in PBS and lysed in RIPA lysis buffer supplemented with cOmplete protease inhibitors on ice for 30 min, followed by centrifugation at 17,900 × *g* for 15 min at 4 °C. Supernatants were mixed with SDS-sample buffer and incubated on ice for 30 min, and analyzed by western blotting.

#### Blue native PAGE

CLPTM1L-KO HEK293 cells expressing CLPTM1L-FH were lysed in DDM lysis buffer (25mM HEPES pH7.4, 150mM NaCl, 1% (19.6 mM) DDM and 1 × protease inhibitor cocktail cOmplete) on ice for 30 min, followed by centrifugation at 17,900 × *g* for 15 min at 4 °C. Supernatants were mixed with NativePAGE sample buffer (Invitrogen) and incubated with various concentrations of SDS for 30 min at RT, resolved using the NativePAGE Bis-Tris Gel System (Invitrogen) and analyzed by western blotting.

### Structure prediction for human CLPTM1L protein

The full sequence of human CLPTM1L (UniProtKB=Q96KA5) was submitted to trRosetta for structure predictions using the default setting (*14*). The trRosetta succeeded in generating CLPTM1L structure models using AtSWEET13 (PDB entry 5XPD) and AtSWEET2 (PDB entry 5CTG) (*18, 48*). The top-ranking model was selected, and the figures were generated by using Open-Source PyMOL 2.3.

### GPI-AP restoration assay in PIGA-KO cells incubated with GlcNAc-PI or GlcN-PI

To measure the steady-state restoration of GPI biosynthesis, PIGA-KO or CLPTM1L-PIGA-DKO HEK293 cells were maintained in full medium in 24-well plate (IWAKI) overnight, and medium were replaced with serum-free DMEM containing GlcNAc-PI or GlcN-PI at various concentrations. Cells were maintained at 37 °C for 24 hours before proceeding to flow cytometry analysis.

To study the time course of the restoration of GPI biosynthesis by GlcNAc-PI or GlcN-PI, PIGA-KO or CLPTM1L-PIGA-DKO HEK293 cells were preincubated in DMEM containing 10 µM GlcNAc-PI or 20 µM GlcN-PI at 37 °C for 2 hours, allowing efficient uptake of GlcNAc-PI or GlcN-PI. Then the cells were washed with serum-free DMEM twice and further incubated in the medium for 0-14 hours, cells harvested at different time points were stained with PE-labeled anti-CD59 and analyzed by flow cytometry. The mean fluorescent intensity values of PIGA-KO HEK293 cells after incubation for 14 hours were taken as 100% restoration.

### Purification of human CLPTM1L-FLAG-6His protein

About 4×10^7^ of Expi293 cells (A14527, Thermo Fisher Scientific) expressing CLPTM1L-FH controlled by Tet on expression system were seeded into two spinner flasks each containing 100 mL of Expi293 expression medium (Thermo Fisher Scientific). Cells were maintained in suspension culture for two and a half days, then cells were cultured with 2 µg/mL doxycycline and 1% FBS for 48 hours to induce high expression of CLPTM1L-FH protein. Cells were collected by spinning at 340 × g for 5 min, and washed twice with 10 mL of cold PBS at 340 × *g* for 5 min. The cell pellets (about 5g) were resuspended in 25 mL of DDM lysis buffer (25mM HEPES pH7.4, 150mM NaCl, 1% (19.6 mM) DDM and 1 × protease inhibitor cocktail cOmplete), and rotated at moderate speed for 1 hour at 4 °C. The cells were centrifuged at 21,900 × *g* for 1 hour at 4 °C. The supernatant was filtered through a 0.45-µm filter. For 200 mL of suspension culture, 1.5 mL of anti-FLAG M2 resin (Sigma Aldrich) equilibrated by wash buffer (25mM HEPES pH7.4, 150mM NaCl, 0.1% DDM) was used. The supernatant was transferred to a new tube containing the anti-FLAG resin and gently mixed on a Nutator for 2 hours at 4 °C. The beads were washed with 10 mL of wash buffer three times per mL of anti-FLAG resin. To elute the protein from the anti-FLAG resin, 1 mL of 500 µg/mL FLAG peptide (F3290, Sigma Aldrich) in FLAGElute buffer per mL of anti-FLAG resin were added. This process was repeated twice each for 1 hour each. The twice eluted samples (8 mL) were combined and kept on ice overnight. Next, the eluted samples were adjusted to equilibration buffer (25mM HEPES pH7.4, 300mM NaCl, 0.1% DDM, 10 mM imidazole). The samples were mixed with 250 µL of HisPur Ni-NTA resin (88221, Thermo Fisher Scientific) equilibrated by 2 mL of equilibration buffer on an end-over-end rotator for 60 min at 4 °C. After centrifugation at 700 × g for 5 min, supernatants were discarded without disturbing the resin pellet, and the resin was washed by 2 mL of equilibration buffer twice, then maintained in 2mL of equilibration buffer. The resin was transferred to a 2mL gravity-flow Poly-prep column (Bio-Rad). The column was washed twice with 2 mL of HisWash buffer (10mM HEPES, pH7.4, 150 mM NaCl, 0.1% DDM, 25 mM imidazole). CLPTM1L-FH protein was eluted 5 times by applying 500 µl of HisElution Buffer (10mM HEPES pH7.4, 100mM NaCl, 0.2% DDM, 250 mM imidazole) to the column, and the column was locked for 3 min. The 500 µl fractions were collected and pooled to 2.5 mL in total. Imidazole in the eluted samples was removed through dialysis using Slide-A-Lyzer Dialysis Cassette MWCO 10K (66380, Thermo Fisher Scientific) in 3L of dialysis buffer (10mM HEPES, pH7.4, 150mM NaCl) for 4 hours, 16 hours, and another one day. CLPTM1L-FH proteins and BSA standards (100ng, 200ng, 500ng, and 1000ng) were separated on SDS-PAGE followed by Imperial Protein Stain (24615, Thermo Fisher Scientific) to quantify the concentration of CLPTM1L-FH. Each tube containing about 32 µg of CLPTM1L-FH were snap-frozen and kept at –80 °C.

### Preparation of [^3^H]GlcN-PI

Lysates of CHO-PA16.1 (PIGU-deficient) cells stably expressing FLAG-PIGL, GST-PIGA, MPDU1-FLAG and GST-DPM2 (*49*) were used expecting efficient synthesis of GlcN-PI. Approximately 1×10^8^ cells were pretreated with 5 µg/mL tunicamycin (Wako) for 3 hours at 37 °C, cells were harvested and washed twice with cold PBS. Cells were then suspended in 4 mL of hypotonic buffer (20 mM HEPES) on ice for 5 min, then mixed with an equal volume of sucrose buffer (20 mM HEPES, 500 mM sucrose). After centrifugation at 600 × *g* for 10 min, the supernatants were transferred to a new tube and spun in a TL-100 ultracentrifuge (Beckman) at 100,000 × *g* for 1 hour. The pellet was suspended in 1mL of reaction buffer (50 mM HEPES-NaOH, pH 7.4, 25 mM KCl, 1 × cOmplete EDTA-free Protease Inhibitor, 1mM GTP, 1mM ATP, 5 mM MgCl_2_, 0.5 mM DTT, 0.2 μg/mL tunicamycin, 20 µCi of 0.2 μM UDP-[6-^3^H]GlcNAc) at 37 °C for 3 hours. The reaction was terminated by adding 3.75 mL of chloroform: methanol (1:2, by volume), and lipids were extracted by the Bligh and Dyer method. The chloroform phase was taken and dried up under N_2_ gas steam. Lipids containing [6-^3^H]GlcN-PI were dissolved in 100 μL of chloroform: methanol (2:1, by volume) and loaded to a high performance thin-layer chromatography (HPTLC) plate (Merck). The [6-^3^H]GlcN-PI was located by phosphorimaging. The [6-^3^H]GlcN-PI were collected by scraping the silica containing [6-^3^H]GlcN-PI off the HPTLC plate and eluted twice with 400 µL of isopropanol/hexane/water (50:25:20, by volume) and sonicated for 20 sec. The lipid solution was collected by centrifugation at 12,000 × *g* and dried up using centrifugal concentrator (TOMY). An aliquot of the purified [6-^3^H]GlcN-PI was analyzed by HPTLC, and the remainder (about 90,000 cpm) was stored at –30 °C.

### Synthesis of C16/C12-NBD-PI

C16/C12-NBD-PI (NBD-PI) was synthesized from C16/C12-NBD-PC (NBD-PC) and myo-inositol by phospholipase D (PLD)-mediated transphosphatidylation as previously described for synthesis of C16/C6-NBD-PI (*24, 50*). An engineered microbial PLD carrying G186T/W187N/Y385R mutations, which can transfer the phosphatidyl group toward the 1-OH group of myo-inositol selectively to synthesize 1-PI, was used (*51*). Briefly, 4 mL of chloroform solution of 1 mg/mL NBD-PC was dried up under N_2_ gas steam and redissolved in 400 µl of ethyl acetate. This lipid solution was mixed vigorously with 320 µL of NaCl-saturated 50 mM acetate buffer (pH 5.6) containing 40 µg of the engineered PLD and 72 mg of myo-inositol for 24 hours at 20 °C. The reaction was terminated by adding 40 µL of 1 M HCl to this biphasic mixture. The lipids were extracted using 800 µL of chloroform: methanol (2:1, by volume). After confirming the formation of the target product by TLC (fig. S5B), the solvent was removed under the N_2_ gas steam. The lipids were redissolved in 500 µL of chloroform and loaded onto a silica gel column (Wakogel C-300, 1 g). NBD-phosphatidic acid (NBD-PA) was eluted with 40 mL of chloroform:methanol (8:2, by volume), and NBD-PI was then eluted with 40 mL of chloroform:methanol (7:3, by volume). Eluents were collected into 2 mL fractions, and analyzed by TLC. The fractions containing NBD-PI were combined and the solvent was removed by a rotary evaporator. The residual lipids were redissolved in 1 mL of chloroform, and loaded again onto a silica gel column (1g) for rechromatography to remove the contaminated NBD-PA. The NBD-PA was eluted with 30 mL of chloroform: petroleum ether: methanol: acetic acid (5:3:1:1, by volume), and NBD-PI was eluted with 70 mL of chloroform: petroleum ether: methanol: acetic acid (4:3:2:1, by volume). The fractions containing NBD-PI were combined, solvent removed by a rotary evaporator with azeotropic removal of acetic acid with toluene, followed by further drying under vacuum to afford NBD-PI (1.5mg, 34%). The purified lipids were analyzed by TLC and Liquid chromatography-mass spectrometry analysis.

### Proteoliposome reconstitution

For the PI-specific phospholipase C (PI-PLC)-based scramblase assay, a previously reported one-pot method was used to make liposomes and proteoliposomes (*21, 25*) using a mixture of lipids including egg PC and brain PS at a molar ratio of 9:1. Briefly, 8.2 μmol egg PC, 0.92 μmol brain PS, and trace amount of [^3^H]PI (≈25,000 cpm) or [^3^H]GlcN-PI (≈12,000 cpm) were combined in a glass screw-cap tube, dried under a stream of the N_2_ gas, and solubilized in 1.9 mL of buffer A (10 mM HEPES-NaOH, pH 7.4,100 mM NaCl, 1% (w/v) TX-100). The lipid solution was vortexed at moderate speed for 20 min until the solution become completely transparent. The solubilized lipids were then mixed with or without purified CLPTM1L-FH in a final volume of 1 mL buffer A, and incubated at 4 °C for 1 hour with end-over-end rotation. For sodium dithionite-based scramblase assays, lipids including egg PC and DOPS at a molar ratio of 9:1 and 0.4 mol% NBD-PC, NBD-PI, or NBD-DOPE were used. Proteoliposomes or protein-free liposomes were generated by two rounds of treatment with 300 mg of wet SM-2 Bio-Beads (Bio-Rad) in total. The samples were placed in 2mL-eppendorf tubes, and end-over-end mixed with 100 mg of wet Bio-beads at RT for 3 hours. A second portion of 200 mg of wet Bio-beads was added and the incubation was continued at 4 °C for 16 hours. Finally, the samples were transferred to 1.5mL-eppendorf tubes without disturbing the Bio-beads and used immediately for scramblase assays.

### PI-PLC-based scramblase activity assay using [^3^H]PI or [^3^H]GlcN-PI

The PI-PLC-based scramblase assay was described previously (*24*). Briefly, 100 μL of liposomes or proteoliposomes containing a trace amount of [^3^H]PI or [^3^H]GlcN-PI were mixed with 10 μL of buffer B (10mM HEPES, pH7.4, 100mM NaCl) and incubated at RT for 5 min. After adding 3 μL of 10 U/mL PI-PLC (P6466, Thermo Fisher Scientific) diluted in buffer B, samples were incubated at 25 °C for 0-10 min. The reaction was stopped by adding 13 μL of 100% ice cold trichloroacetic acid (TCA), then mixed with 5 μL of 4 mg/mL cytochrome c from horse heart (105201, Sigma-Aldrich). The mixtures were kept on ice for 1 hour with occasional vortex at high speed, then centrifuged at 21,900 × *g* for 15 min to precipitate proteins and phospholipids. The resulting supernatant (130 μL) was mixed with 10 mL of Clear-sol II scintillation fluid (09136-83, Nacalai Tesque) and counted with a liquid scintillation counter (TRI-CARB 4810TR, PerkinElmer). TCA samples prepared from samples without PI-PLC were taken as background, and TCA samples taken after PI-PLC treatment for 10 min in the presence of 0.91% Triton X-100 were taken as 100% hydrolysis control.

### Dithionite-based scramblase activity assay using NBD-labeled phospholipids (PLs)

The dithionite-based scramblase assay was described previously (*52*). Briefly, 50 μL of NBD-PL containing liposomes were diluted into 2 mL of buffer B in a disposable fluorescence cuvette (67.754.00002, SARSTEDT) at RT. The total fluorescence intensity was monitored over time using the Hitachi F-2700 fluorescence spectrophotometer (λex = 470 nm, λem = 530 nm). After obtaining a stable fluorescence signal, 40 μL of freshly prepared 1M sodium dithionite (TCI) was added to the cuvette, the fluorescence signal change was monitored for 10 min. Data were collected and analyzed using the FL Solutions 4.1 software (Hitachi).

### In vivo metabolic labeling

To detect mannose-containing GPI intermediates, approximately 2 × 10^6^ of CHO-PIGS-KO, HEK293-PIGS-KO, PIGS-SYVN1-DKO and PIGS-SYVN1-CLPTM1L-TKO cells expressing empty vector, or CLPTM1L were precultured in complete medium overnight. Cells were washed with wash medium (glucose-free DMEM buffered with 20 mM HEPES), then incubated for 1 hour at 37 °C in 1 mL of reaction medium A (glucose-free DMEM buffered with 20 mM HEPES, pH 7.4, and 10 μg/mL tunicamycin (Wako), and 100 μg/mL glucose) supplemented with 10% dialyzed FBS (Gibco). The cells were then incubated in the reaction medium A containing 40 μCi/mL D-[2-^3^H]mannose (American Radiolabeled Chemicals) for 1 hour at 37 °C in 5% CO_2_. To detect GlcN-PI or GlcN-(acyl)-PI, the cells were washed with inositol-free DMEM medium, and incubated in 1 mL of reaction medium B (inositol-free DMEM buffered with 20 mM HEPES, pH 7.4) supplemented with 10% dialyzed FBS in the presence of 10 μCi of myo-[2-^3^H]inositol (PerkinElmer) for 10 hours. After metabolic labeling, the cells were washed twice with 1 mL of cold PBS and pelleted by centrifugation. Lipids and radiolabeled GPIs were extracted by 1-butanol (Wako) partitioning, separated by HPTLC (Merck), and visualized by a FLA 7000 analyzer (Fujifilm). Quantitative analysis was performed by JustTLC (SWEDAY).

